# O_2_ exchange profiling reveals physiological diversity among taxonomically characterized green microalgal strains

**DOI:** 10.64898/2026.11.23.768518

**Authors:** Michael Kikoine, Tamar Sagi, Shahar Malina, Daniel Paldi, Liat Oren-Young, Alla Alster, Ruth N. Kaplan-Levy, Iftach Yacoby

## Abstract

Taxonomic identification provides essential information about microalgae, but a shared species assignment does not necessarily imply similar physiological behavior. This study combined microscopy, ribosomal DNA sequencing, and O_2_ exchange measurements to characterize 20 green microalgal strains, comprising 18 environment-derived and two previously identified model strains. Net O_2_ evolution and post illumination consumption were measured across 17 increasing irradiances in three independently grown cultures per strain. Maximum differences between pair means for net O_2_ evolution were 18.12, 7.82, and 10.50 µmol O_2_ L⁻¹ min⁻¹ for the *Chlorella sorokiniana*, *Desmodesmus abundans*, and *Tetradesmus*/*Acutodesmus obliquus* pairs at 100, 50, and 300 µmol photons m⁻² s⁻¹, respectively; all three cultures of one strain exceeded all three of the other at those irradiances. Within-pair ITS sequences differed by at most one nucleotide over the shared regions. Separately normalized profiles formed five exploratory groups, but group membership was sensitive to culture resampling, the number of principal components, and rate-extraction windows. Physiological distances showed no clear association with 18S patristic distances or assigned genus. Interpretation is limited by provisional species assignments, three cultures per strain, and a single ascending irradiance sequence that confounds light intensity with elapsed time and unmeasured carbon availability. O_2_ exchange profiling therefore provides complementary information about individual strains under a common experimental protocol, without establishing stable physiological classes or steady-state capacities.

## Introduction

Accurate identification of green microalgae can be challenging because of similar phenotypes occur in distinct evolutionary lineages, while diagnostic morphological characters may vary with environmental conditions. This difficulty is particularly apparent among small coccoid algae, which offer relatively few features that can be distinguished by light microscopy. Studies combining morphological observations with molecular analyses have revealed diversity that would be difficult to recognize from appearance alone, supporting an integrated approach to microalgal identification (Fawley et al., 2004; Bock et al., 2011; Darienko et al., 2015). Establishing taxonomic identity nevertheless addresses only part of the characterization of an algal strain; its physiological responses must also be examined to understand its functional properties (Kessler & Huss, 1992).

Ribosomal DNA sequencing provides an important complement feature to microscopy by placing morphologically similar organisms within a molecular framework. Analyses combining the small-subunit ribosomal region with the more variable internal transcribed spacers have helped distinguish closely related green algae and revise classifications based on morphology (Bock et al., 2011). In Lake Kinneret phytoplankton, integrating morphological identification with sequencing of the *SSU–ITS* region and *rbcL* supported existing identifications, clarified uncertain assignments, and enabled taxonomic updates (Kaplan-Levy et al., 2016). Such approaches strengthen identification, but their resolution depends on the information contained in the selected markers and the available reference sequences (Zou et al., 2016). They do not establish equivalence across the remainder of the genome or directly describe physiological behavior.

The distinction between taxonomic assignment and functional characterization is relevant even among strains carrying the same species name. Comparative genomic analysis of three strains assigned to *Chlorella sorokiniana* revealed substantial differences in genome content, illustrating that a shared designation does not establish genomic equivalence (Hovde et al., 2018). Physiological studies likewise demonstrate the value of examining individual strains. The desert biological soil crust alga *Chlorella ohadii*, although closely related to *C. sorokiniana* in molecular analyses, exhibited distinctive structural and photosynthetic characteristics, including strong resistance to excess irradiance and responses that depended on the carbon source supplied during growth (Treves et al., 2013). A later genome comparison identified features potentially associated with rapid growth and resistance to photodamage in this alga, including additional ribosome-encoding genes and candidate genes involved in metabolic flexibility (Murik et al., 2024). These findings motivate direct physiological comparisons among identified strains while emphasizing the importance of the conditions under which their responses are measured.

O_2_ exchange measurements provide a means of characterizing responses associated with photosynthesis and respiration. Measuring net O_2_ evolution during illumination and O_2_ consumption during subsequent darkness across an irradiance series produces response profiles that contain more information than a single rate measurement. However, these profiles depend on the experimental sequence as well as the irradiance applied. In *Chlamydomonas reinhardtii*, respiratory O_2_ consumption following illumination increased with preceding light exposure, demonstrating that post illumination respiration can retain the influence of recent photosynthetic activity (Xue et al., 1996). Comparisons among strains therefore require a defined cultivation and measurement protocol, with the resulting profiles interpreted as responses under those conditions.

Absolute rate magnitudes and relative response patterns describe different aspects of physiological variation. Normalizing evolution and respiration separately within each strain emphasizes how each process changes across irradiance relative to its own strongest measured response. This approach facilitates comparison of response patterns while removing differences in peak magnitude and the absolute scale of respiration relative to evolution. Hierarchical clustering and principal component analysis can summarize similarities and variation among these multivariable profiles, while k-means clustering can provide an exploratory grouping of strains. Comparing these physiological groupings with morphological observations and molecular relationships offers a way to examine the additional information obtained from functional measurements. Physiological differences under the tested conditions can inform strain characterization, but their taxonomic significance requires separate evidence. In the present study, we combined microscopy, rDNA-based identification, and physiological measurements to characterize a collection of 20 green microalgal strains, including environmental isolates and previously characterized strains of *Chlorella ohadii* and *Acutodesmus obliquus*. Cellular morphology and dimensions were examined alongside ribosomal DNA sequences, and O_2_ evolution and respiration were measured during a standardized sequence of alternating light and dark intervals across increasing irradiance. Normalized response profiles were compared using hierarchical clustering, PCA, and k-means clustering, paired evolution–respiration trajectories were used to interpret variation within the resulting groups, and the adjusted Rand index quantified agreement between the two physiological groupings. Mantel and PERMANOVA analyses evaluated correspondence with *18S rRNA* phylogenetic distances and taxonomic assignments, respectively. The objective was to determine what additional functional information these measurements provided and to explore the correspondence between physiological similarity, morphological resemblance, and assigned taxonomic identity within the collection. Particular attention was given to strains assigned to the same species or displaying similar morphology.

## Methods and Materials

### Algal strains

The algal strains analysed in this study originated from two sources. Reference strains, that were maintained in the lab, isolates obtained from the Israel National Culture Collection of Algae (INCCA)(Kaplan-Levy et al, 2020), and additional strains isolated and established in culture by our laboratory. Following characterization, the newly isolated strains were deposited in the INCCA for long-term safekeeping and future access. The origin and accession numbers of all strains are provided in Table S1.

### Reference Culture sources

The reference culture G1102, supplied as *Acutodesmus obliquus*, was obtained from the Culture Collection of Algae at Göttingen University (SAG 276-3d) and was originally isolated in Italy in 1954. The supplied taxonomic name was retained throughout the manuscript. Strain G1093 originates from the biological sand crust at the Nitzana (Nizzana) dunes in the north-western Negev and is identified as *Chlorella ohadii*. The laboratory-held G1093 culture corresponds to the culture catalogued as SAG 2657, according to laboratory provenance; the culture used here was not obtained directly from SAG. This name was validated by Treves and Guiry (2026), who designated type material collected at the same locality in 2012; the organism had previously been referred to as *C. sorokiniana* subsp. *ohadii* (Murik et al., 2024).

### Strain isolation (this study)

#### Sampling

Water samples (50 mL) were collected from the sites shown in Figure 1 into sterile 50-mL tubes. At each sampling site, 5-mL aliquots were inoculated onto Tris–phosphate (TP) agar plates, prepared by omitting acetate from Tris–acetate–phosphate (TAP) medium (Gorman & Levine, 1965). Plates were either antibiotic-free or supplemented with carbenicillin at 100 µg mL⁻¹. An additional 5-mL aliquot was transferred to a flask containing 50 mL of liquid TP medium. Cultures were incubated for 12–48 h, depending on algal growth, at 20–25 °C under continuous illumination of 100–200 µmol photons m⁻² s⁻¹. Plates were maintained on a rack, whereas liquid cultures were incubated with shaking at 120–150 rpm.

**Figure 1:**
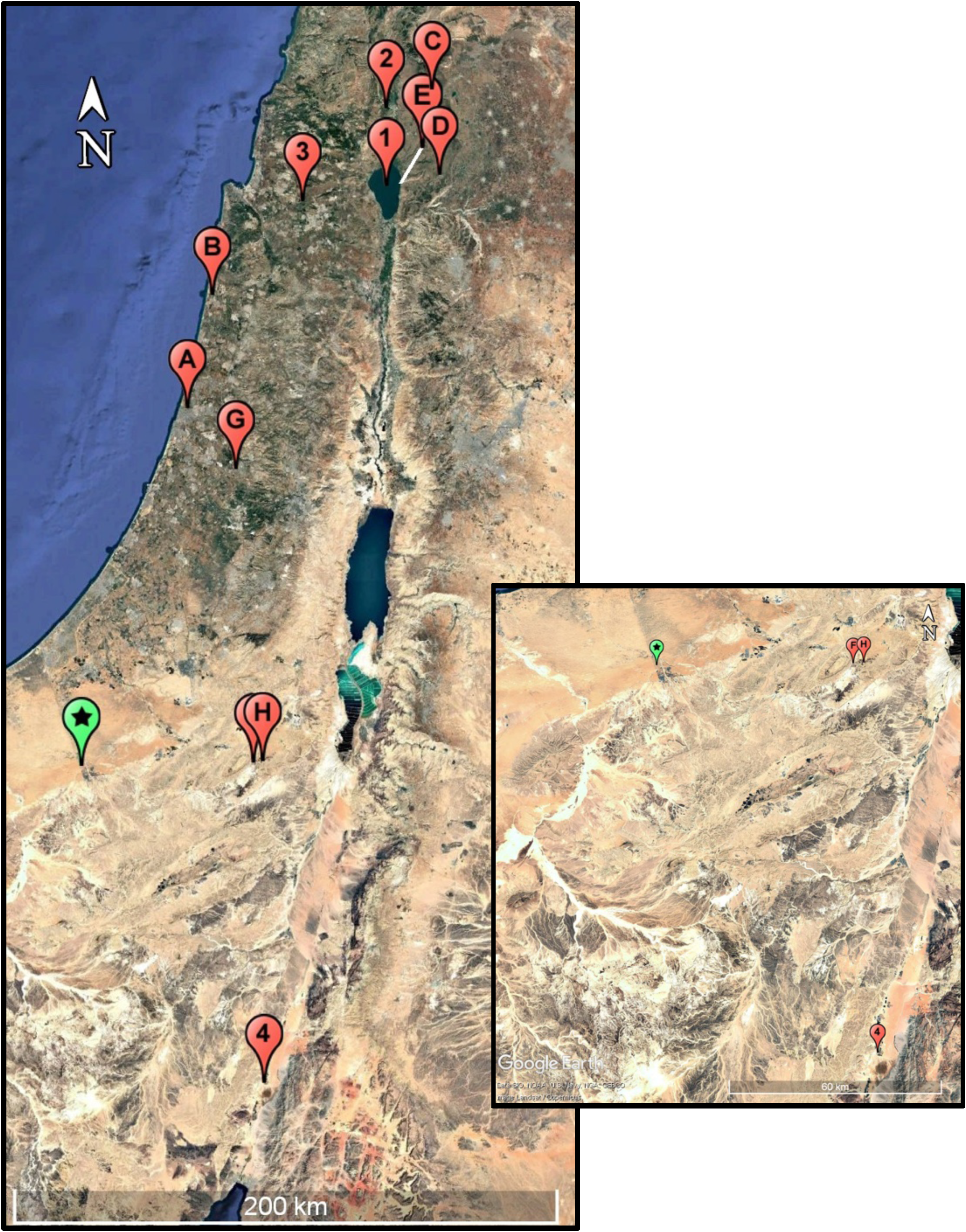
Satellite image of the locations with inset on the Negev region. 12 of the 20 tested strains were obtained at: (A) Yarkon River (B) Hadera Stream (C) Bental Reservoir (D) El-Al Stream (E) Kursi Beach (F) Ein Yorqe’am (G) Modi’in (H) Gev Yamin 7 of the 20 tested strains purchased from INCCA: (1) Lake Kinneret – Station A (2) ake Hula (3) Eshkol Reservoir (4) Yotveta Reservoir (Green Star) Nitzana Dunes – *Chlorella ohadii*. Exact coordinates for each location are provided in Table S1. Satellite image and pinpoints were generated using Google Earth.

### Single-cell isolation and culture establishment

Individual algal cells were isolated by micromanipulation using a SporePlay instrument (Singer Instruments) on antibiotic-free Tris–acetate–phosphate (TAP) agar containing 0.1% (v/v) glacial acetic acid. Cells were dragged across the agar surface to spatially separated positions and incubated until visible colonies developed, requiring 3 days to 3 weeks depending on the isolate (Figure 2). Plates were visually inspected during colony development, and colonies from plates showing no visible bacterial or fungal growth were selected for subsequent cultivation. Axenicity was not tested. Colony selection was based on visual inspection only; the cultures are therefore regarded as unialgal but not necessarily axenic, and a bacterial contribution to O_2_ consumption cannot be excluded.

**Figure 2:**
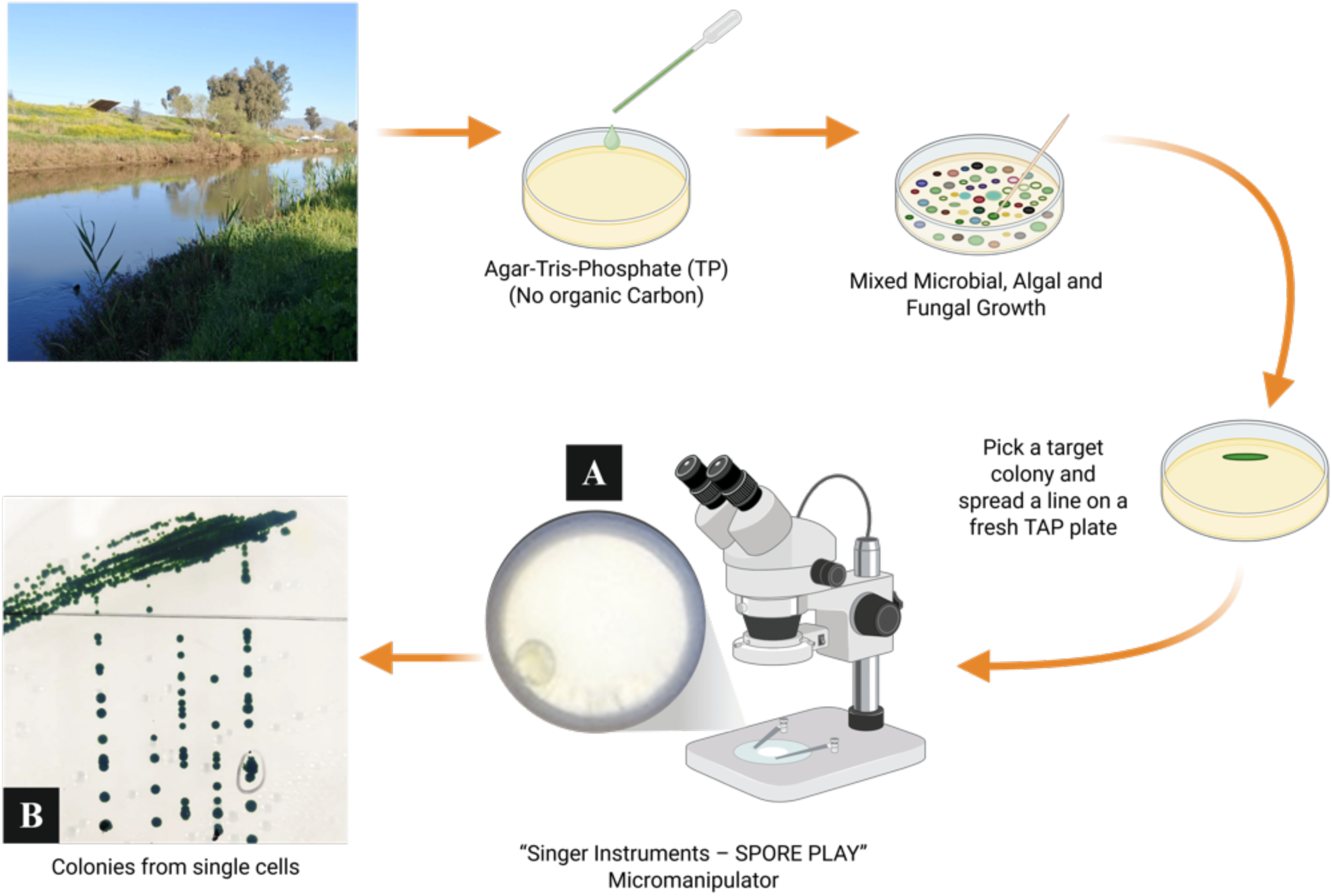
Workflow for environmental sampling and single-cell isolation of microalgae. Water samples were inoculated onto Tris–phosphate (TP) agar to establish mixed cultures. Material from selected colonies was streaked onto antibiotic-free Tris–acetate–phosphate (TAP) agar containing 0.1% (v/v) glacial acetic acid. Individual algal cells were then isolated using a SporePlay micromanipulator (Singer Instruments). Multiple cells from the same sample were moved to spatially separated positions on the same plate. (A) Microscope image of the glass micromanipulation needle tip carrying an individual algal cell; the circular structure is the needle tip. (B) Microalgal colonies 21 days after micromanipulation, arranged along the paths used to separate and deposit individual cells.

### Microscopy and cell morphometry

Live, unfixed algal cells were imaged by brightfield microscopy using an EVOS M5000 microscope equipped with Olympus 60× (numerical aperture, NA = 1.42) and 100× (NA = 1.45) oil-immersion objectives, with images recorded in RGB format. More than 50 cells per isolate were measured from a single young culture with a total chlorophyll concentration below 20 µg mL⁻¹. All cells in focus within the acquired images were included in the analysis. Cell dimensions were quantified using custom image-analysis software, Algal Detector v1.5.1. The accompanying data and code package includes the Algal Detector v1.5.1 research release and source snapshot (https://github.com/energylabtau-alga/algal-detector/releases/tag/v1.5.1); the version used for each historical measurement was not recorded. Pixel-to-micrometer conversion was calibrated separately for each objective using scale bars generated by the microscope imaging software, with a consistent spatial scale maintained across images acquired at the same magnification. Cell length was defined as the longest straight-line segment contained within the cell outline, and width as the longest internal segment perpendicular to the length axis. Cell outlines were superimposed separately for each strain to visualize typical cell shape and size and variation among the measured cells. Representative micrographs were selected to illustrate typical cellular morphology (Figure S1). Morphological identification was supported by comparison with reference images from the Culture Collection of Algae and Protozoa (CCAP) and the Culture Collection of Algae at Göttingen University (SAG), together with morphological descriptions available through AlgaeBase (Guiry & Guiry, 2026). Taxonomic assignments of the environmental isolates are summarized in Table S1 and discussed in the legend to Figure 4.

**Table 1:** Working taxonomic assignments of the 20 Chlorophyte strains, based on rDNA sequence analysis, morphology, and culture provenance. Strains are ordered by INCCA-code. Panel identifiers refer to Figure 4 and link to collection sites in Figure 1: letters A–H indicate strains isolated in our laboratory, numbers 1–3 indicate strains obtained from INCCA, and the green star indicates the Nitzana Dunes origin of G1093 (*Chlorella ohadii*). Additional numeric suffixes distinguish strains from the same site. G1102 (*Acutodesmus obliquus*; SAG 276-3d) was purchased from SAG and has no sampling site in Figure 1. G1093 and G1102 were included as previously identified model strains. *Acutodesmus obliquus*, the supplied name retained for G1102, is a synonym of *Tetradesmus obliquus*, the assignment used for G1097. “sp.” indicates a genus-level name and “cf.” a tentative species assignment. AlgaeBase identifiers refer to species records or, for “sp.” entries, genus records. Species names remain working assignments; molecular evidence and unresolved discrepancies are detailed in Table S6.

| **Index** | **Panel Identifier (Figure 4)** | **INCCA Code** | **Species Identification** | **AlgaeBase ID** |
|---|---|---|---|---|
| **1** | 1.1 | G1014 | *Chlamydomonas* sp. | 43319 |
| **2** | 1.2 | G1044 | *Pediastrum duplex* | 27511 |
| **3** | 1.3 | G1049 | *Desmodesmus denticulatus* | 42405 |
| **4** | 1.4 | G1058 | *Haematococcus lacustris* | 36266 |
| **5** | 1.5 | G1070 | *Stauridium tetras* | 129243 |
| **6** | 2 | G1074 | *Coelastrella* sp. | 45299 |
| **7** | Green Star | G1093 | *Chlorella ohadii* | 177095 |
| **8** | A1 | G1094 | *Chlorella sorokiniana* | 51508 |
| **9** | A2 | G1095 | *Micractinium tetrahymenae* | 176401 |
| **10** | E | G1096 | *Desmodesmus abundans* | 42384 |
| **11** | F1 | G1097 | *Tetradesmus obliquus* | 158061 |
| **12** | F2 | G1101 | *Chlorolobion braunii* | 27677 |
| **13** | SAG | G1102 | *Acutodesmus obliquus* | 135513 |
| **14** | D | G1103 | *Lobochlamys segnis* | 118596 |
| **15** | A3 | G1104 | *Monoraphidium cf. dybowskii* | 131950 |
| **16** | B | G1105 | *Micractinium* sp. | 43421 |
| **17** | C | G1107 | *Desmodesmus abundans* | 42384 |
| **18** | 3 | G1114 | *Micractinium* sp. | 43421 |
| **19** | G | G1121 | *Chlorella lewinii* | 147234 |
| **20** | H | G1123 | *Chlorella sorokiniana* | 51508 |

### Molecular identification

Genomic DNA was extracted using a ZymoBIOMICS DNA Miniprep Kit (Zymo Research cat # D4300). A nuclear ribosomal DNA region spanning the *18S rRNA* gene, *ITS1*, the *5.8S rRNA* gene, *ITS2*, and part of the *28S rRNA* gene was amplified using primers 1 and 2 (White et al., 1990, Table S2), following Kaplan-Levy et al. (2016). PCR was performed using VeriFi Hot Start Mix (PCR Biosystems), with the reaction composition and thermal cycling conditions provided in Table S3. Amplification products were examined by electrophoresis on a 1% (w/v) agarose gel. Representative products from G1058 and G1044 migrated between the 2 and 3 kb markers (Figure S2). Because G1058 yielded multiple bands, the selected band was excised and purified using the NucleoSpin Gel and PCR Clean-up Mini kit (MACHEREY-NAGEL, cat # 740609.250) before Sanger sequencing; the same kit was used to purify PCR products that did not require gel separation. Products were prepared for Sanger sequencing according to the sequencing facility’s requirements (10 ng DNA per 100 nucleotides for PCR product and 10 pmol primer in total). Standard Sanger sequencing used primers 1–4 (Figure 3, Table S2). Reads were edited and assembled in BioEdit (Hall, 1999), with reverse reads reverse-complemented for assembly. Consensus sequences were initially compared with reference sequences using NCBI Nucleotide BLAST. Working taxonomic assignments integrated cell morphology, culture provenance, and similarity to reference sequences associated with CCAP, SAG, or KLL/INCCA (Kinneret Limnological Institute, Israel Oceanographic & Limnological Research/Israel National Culture Collection of Algae) strains. The ≥98% values reported for matches to the original reference sequences indicate sequence identity, not query coverage. Neither high sequence identity nor high query coverage was considered sufficient to establish species identity. The 20 study contigs were also compared locally with the 127 public reference sequences retained for the 18S phylogeny using archived NCBI BLAST+ megablast output (Camacho et al., 2009), after confirming that the retained query and reference sequences were unchanged. Strongest-high-scoring segment pair (HSP) identity and coverage were distinguished from total query coverage (Table S6). ITS1 and ITS2 were delimited by transfer of public reference annotations using Biopython (Cock et al., 2009) and compared only over regions present in both members of each nominal species pair (Table S7; Supplementary Methods). These analyses did not constitute a new database-wide search or an ITS2 secondary-structure analysis. Reference names and sequence similarity were not treated as proof of species identity; unresolved discrepancies are reported explicitly. Study-sequence accession information is provided in Table S6.

**Figure 3:**
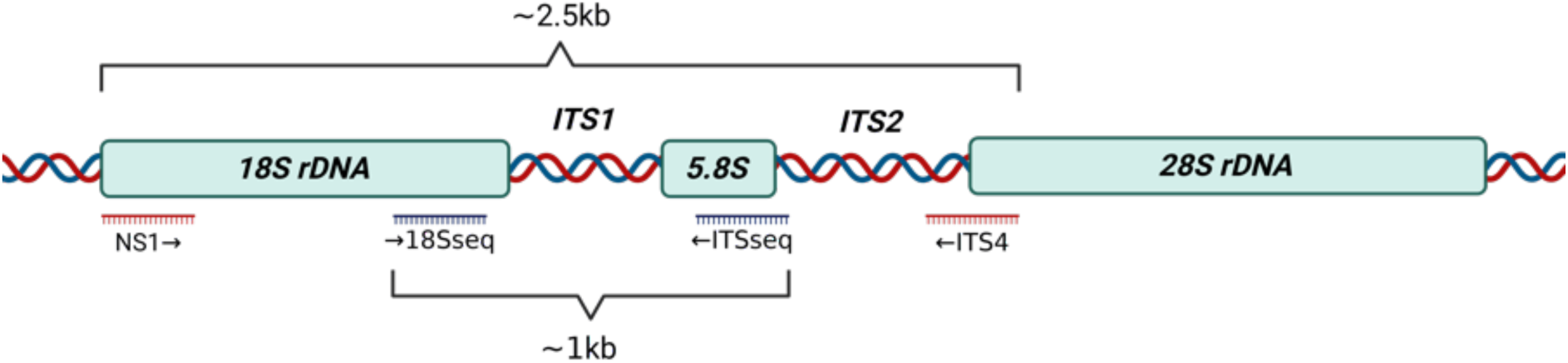
Schematic representation of the ribosomal DNA region and primer arrangement used for molecular identification of green microalgae. Primers 1 and 2 were used to amplify a region spanning the *18S rRNA* gene, *ITS1*, the *5.8S rRNA* gene, *ITS2*, and part of the *28S rRNA* gene. Sanger sequencing was performed using primers 1–4. Primer positions and directions of sequence extension are indicated. Primer sequences are provided in Table S2, and PCR reaction composition and thermal cycling conditions are provided in Table S3. The figure is not drawn to scale.

**Figure 4:**
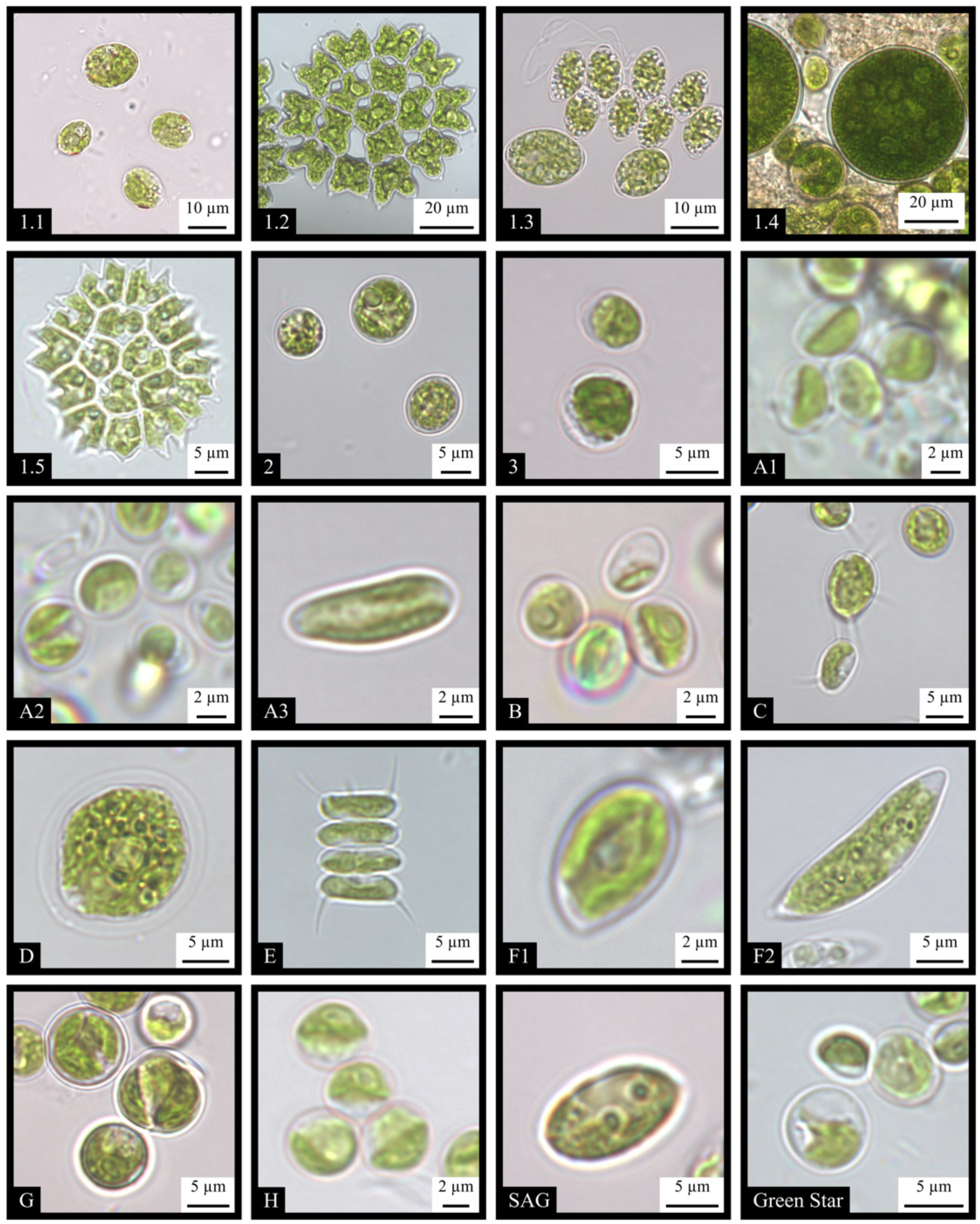
Representative bright-field micrographs of live cells and colonies of the 20 green microalgal strains used in this study, acquired using an EVOS M5000 microscope equipped with an Olympus 100× oil-immersion objective (NA 1.45). Panel identifiers correspond to the strain entries in Table 1, with collection sites shown in Figure 1. Panels SAG (G1102, *Acutodesmus obliquus*) and “Green Star” (G1093, *Chlorella ohadii*) represent previously identified model strains. Scale bars are indicated in each panel.

Ribosomal DNA (rDNA) contigs from the 20 green microalgal strains, phylum Chlorophyta, were aligned using MUSCLE (Edgar, 2004), implemented in MEGA12 (Kumar et al., 2024). Sequences from G1105 and G1123 were reverse-complemented before alignment to standardize their orientation. For the main phylogenetic analysis (Figure 5), the study sequences were aligned with 127 database reference sequences and the outgroup sequence *Pedinomonas tuberculata* HE610134.1, giving 148 sequences in total. The alignment was trimmed at the 18S–ITS1 boundary. Partial deletion at 95% site coverage retained 1,656 aligned 18S positions. Maximum-likelihood inference used the Tamura–Nei nucleotide-substitution model with gamma-distributed rate variation and invariant sites (TN93+G+I), selected according to the lowest Bayesian information criterion score in MEGA12 (Precise model elaborated in the legend of figure 5). The resulting tree was rooted using *P. tuberculata*. A separate analysis restricted to the rDNA contigs of the 20 study strains is presented in Figure S6. This analysis used the Tamura–Nei model with gamma-distributed rate variation and invariant sites (TN93+G+I). Partial deletion at 95% site coverage retained 2,130 alignment positions. No outgroup was included in this analysis, and its topology was therefore interpreted as unrooted. In both analyses, gamma-distributed rate variation was approximated using five discrete categories, tree searches used nearest-neighbor interchange starting from automatically generated initial trees, and branch support was evaluated using 1,000 standard bootstrap replicates. Branch lengths were estimated in nucleotide substitutions per site.

**Figure 5:**
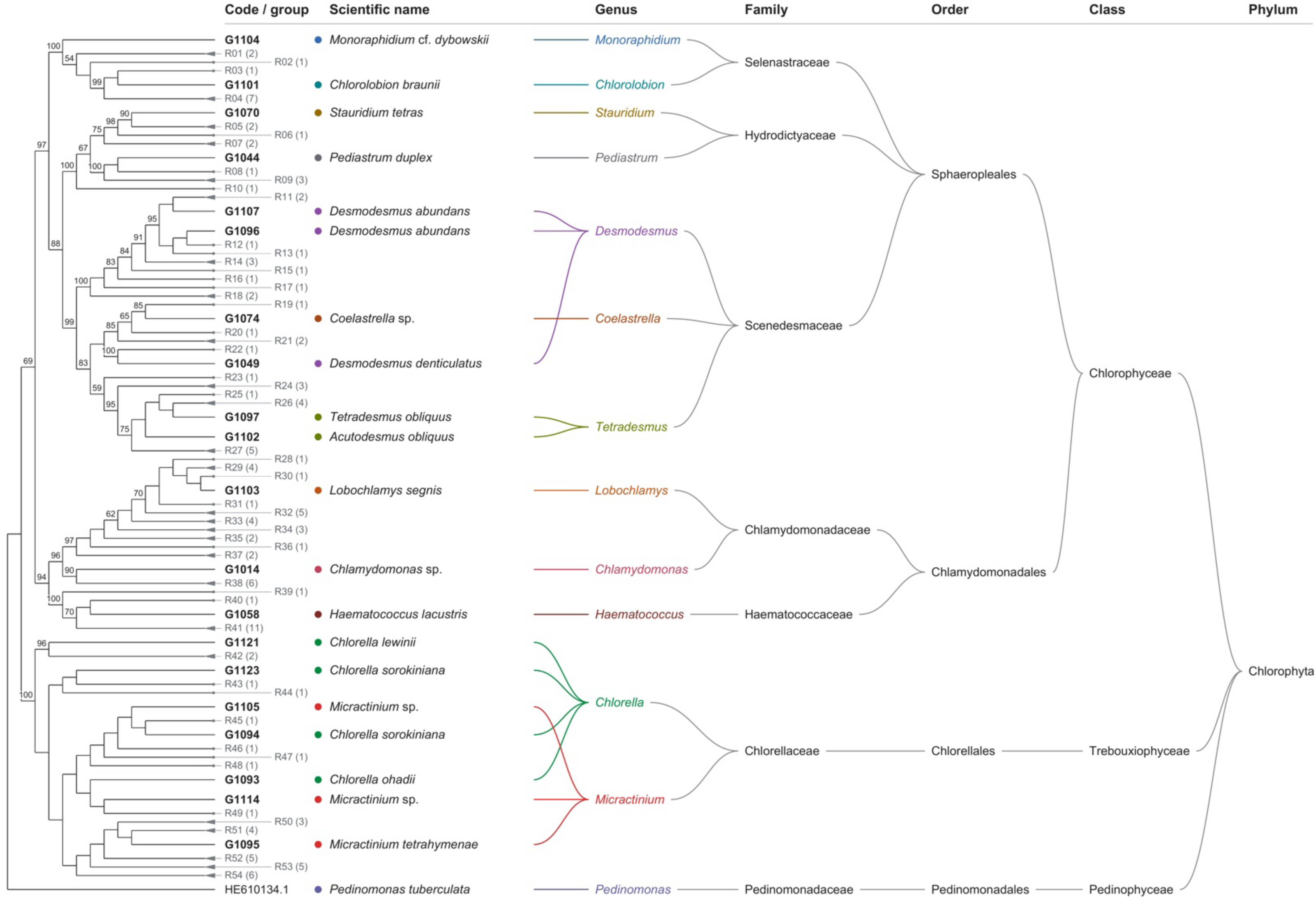
Rooted *18S rRNA* gene phylogeny and taxonomic classification of the 20 microalgal strains used in this study. The maximum-likelihood tree was inferred in MEGA12 from 148 sequences comprising 20 study strains, 127 database references, and *Pedinomonas tuberculata* (HE610134.1) as the outgroup. Analysis used the Tamura–Nei model with gamma-distributed rate variation among sites (five categories) and invariant sites (TN93+G+I). Partial deletion at 95% site coverage retained 1,656 aligned 18S positions. Branch numbers indicate bootstrap support from 1,000 replicates, rounded to the nearest integer; values below 50%, labels on zero-length branches, and overlapping labels are omitted. Branch spacing is schematic and does not represent evolutionary distance. Study strain codes are bold. All reference sequences are represented by 54 reference-only branches labelled R01–R54, with sequence counts in parentheses. Triangles denote collapsed clades containing multiple references; branches with a count of one represent individual reference sequences. The complete tree is shown in Figure S7. Colored circles and connecting lines associate the displayed scientific names with their assigned genera. The remaining columns show family, order, class, and phylum. These columns are taxonomic annotations, rather than an independently inferred phylogeny. Study names retain the identifications reported in Table 1. The name *Acutodesmus obliquus* is retained for G1102 and linked to *Tetradesmus* in the genus column to reflect synonymy. The accession key is provided in Table S4.

For visualization, branches were displayed schematically, and selected nodes were rotated to improve label arrangement without changing the inferred relationships. In Figure 5, reference-only clades were summarized as 54 groups (R01–R54), while all study strains and the outgroup remained individually labelled. The fully expanded tree is provided in Figure S7, and the accession numbers and scientific names of all 127 references are listed in Table S4. Accompanying taxonomic annotations summarize the classification associated with the assigned strain identities from genus to phylum; these annotations do not represent a separately inferred phylogeny.

### Chlorophyll quantification

Total chlorophyll was measured by dimethyl sulfoxide (DMSO) extraction. A 1-mL culture sample was centrifuged at 16,000 × g for 1 min, resuspended in 1 mL DMSO, and incubated at 65°C for 15–20 min. After a second centrifugation at 16,000 × g for 1 min, extract absorbance was measured at 665, 649, and 730 nm. Total chlorophyll concentration (µg mL⁻¹) was calculated as 12.19(A₆₆₅ − A₇₃₀) − 3.45(A₆₄₉ − A₇₃₀) + 21.99(A₆₄₉ − A₇₃₀) − 5.32(A₆₆₅ − A₇₃₀), summing chlorophyll a and b according to Wellburn (1994). The 730-nm reading provided a baseline correction.

### O_2_ kinetics measurements

Algal cultures were grown in 150 mL flasks fitted with gas-permeable sponge caps and containing 50 mL of BG-11 medium supplemented with a vitamin mix (Supplementary Methods). Cultures were maintained at 25°C with shaking at 150 rpm under continuous irradiance of approximately 100–200 µmol photons m⁻² s⁻¹ in a CO_2_-controlled incubator (Multitron, Infors HT). The incubator atmosphere was maintained at 2% CO_2_ using a probe that regulated delivery from a cylinder containing 100% CO_2_. CO_2_ enrichment was used during cultivation to accelerate biomass accumulation. Cultures were harvested at a total chlorophyll concentration below 20 µg mL⁻¹, centrifuged, and resuspended in TAP medium at 20 µg total chlorophyll mL⁻¹. No bicarbonate was added. The pH, dissolved inorganic carbon concentration, and cell counts were not measured. A 1mL aliquot was transferred to an ALGi measurement chamber equipped with a fiber-optic O_2_ sensor connected to a FireSting-O_2_ meter (PyroScience). Chamber temperature was maintained at 25°C using a circulating water system. Samples were continuously mixed using miniature magnetic stir bars, with the same stir bars and stirring speed across measurements. No CO_2_ was supplied during the assay. White-light illumination was controlled using ALGi software, and dissolved O_2_ concentrations were recorded using Pyro Oxygen Logger version 3.213. After an initial 1min dark interval, samples were exposed to 17 alternating 1min light and post illumination dark intervals (Figure S3), with irradiance increasing sequentially through 50, 100, 150, 200, 250, 300, 350, 400, 450, 500, 550, 600, 650, 700, 800, 1000, and 2000 µmol photons m⁻² s⁻¹. The approximately 35 min series always used the same irradiance order, confounding irradiance with elapsed time, accumulated dissolved O_2_, and possible depletion of inorganic carbon. Rates were estimated by linear regression over candidate 30 sec windows beginning between 15 sec before and 15 sec after each nominal light or dark transition, at 1.2 sec increments. The maximum slope was selected for light-associated O_2_ evolution and the minimum for dark-associated O_2_ consumption (Figures S3–S5). Candidate windows could therefore overlap a transition. Light-associated measurements represented net O_2_ evolution without correction for respiratory consumption; dark-associated measurements represented post illumination O_2_ consumption, which can differ from respiration after prolonged dark adaptation (Kliphuis et al., 2011). The final analysis included three independently grown cultures per strain, giving 60 biological replicates across 20 strains. A sensitivity analysis re-extracted all rates from the fixed central 15–45 s of each nominal interval using the same interpolation and regression procedure (Figure S10).

### Data processing and hierarchical clustering

For each strain and irradiance level, O_2_ evolution and respiration rates were averaged across the three retained biological replicates before normalization. Mean profiles were normalized separately for each strain and process across the 17 irradiance levels. Evolution rates were divided by the maximum mean evolution rate, whereas respiration rates were divided by the most negative mean respiration rate, yielding dimensionless values in which 1 represented the strongest response within each process. Negative normalized values indicated rates opposite in sign to the corresponding normalization reference. Figure 6 displayed these normalized values, with strains represented by rows and irradiance levels by columns. The 17 evolution and 17 respiration values were concatenated into one 34 variable vector per strain. All 20 vectors were complete, so no missing-value imputation was required. Agglomerative hierarchical clustering was performed in SciPy using correlation distance, defined as 1 − Pearson’s correlation coefficient, and average linkage. Intercluster distances were calculated as the means of all pairwise distances between strains belonging to the two clusters. Optimal leaf ordering was applied for visualization without changing the clustering hierarchy. Internal node labels were calculated as 1 minus the corresponding linkage distance, equivalent to the mean pairwise Pearson correlation between the two clusters being joined. Dendrogram branches were displayed schematically rather than in proportion to clustering distance.

**Figure 6:**
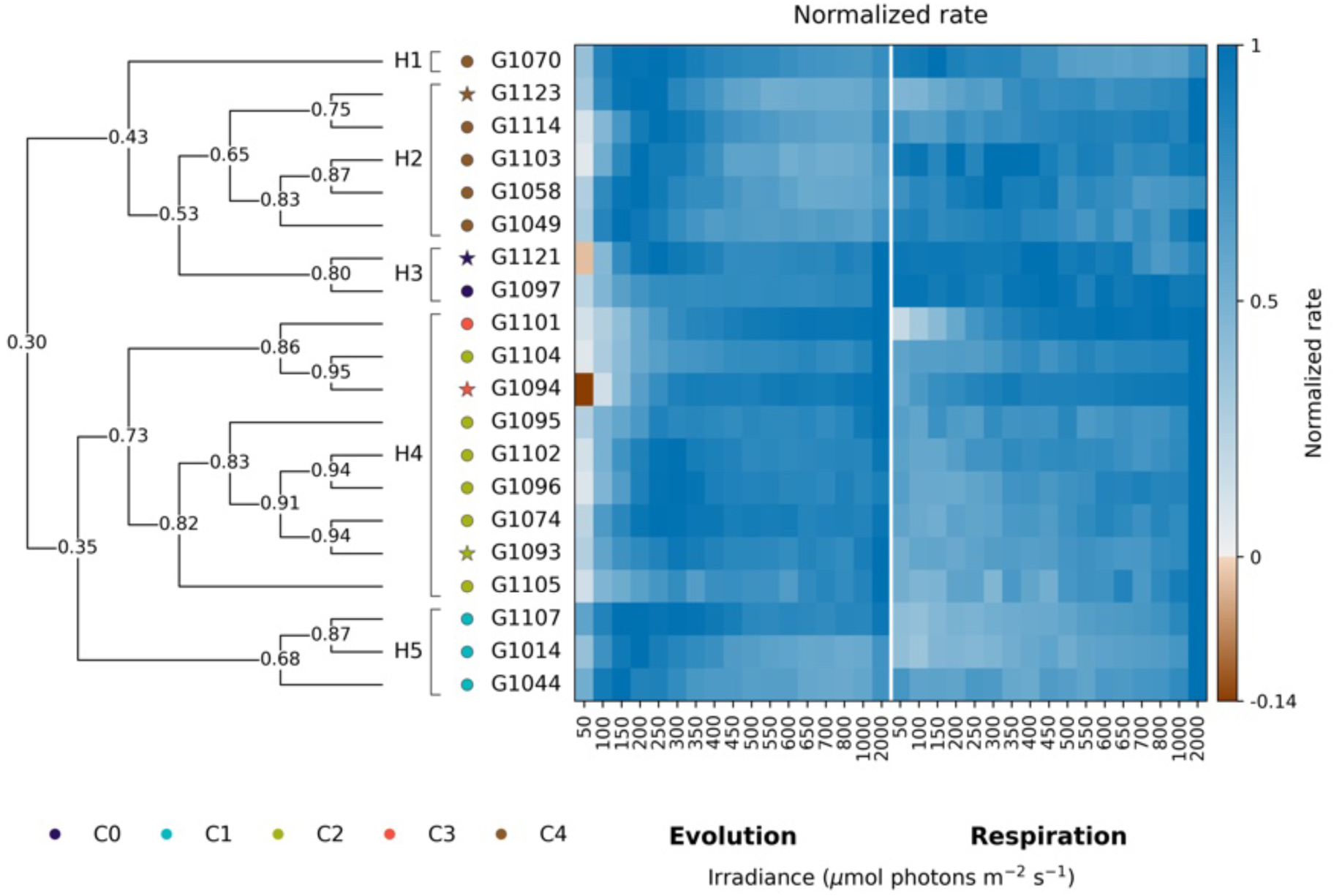
Heatmap and hierarchical clustering of 20 green microalgal strains based on normalized O_2_ evolution and respiration profiles. Rows represent strains, and columns represent 17 irradiance levels spanning 50–2,000 µmol photons m⁻² s⁻¹ for evolution (left) and respiration (right), separated by a white line. Rates were averaged across biological replicates before normalization. For each strain, evolution rates were divided by the maximum mean evolution rate and respiration rates by the most negative mean respiration rate, yielding a value of 1 for the strongest response within each process. Blue indicates positive normalized values, white indicates zero, and brown indicates values opposite in sign to the normalization reference. Each strain’s two profiles were concatenated into a 34-variable vector and clustered using correlation distance (1 − Pearson’s correlation coefficient) and average linkage, with optimal leaf ordering applied for display. Numbers at internal nodes indicate the mean pairwise Pearson correlation between strains belonging to the two groups joined at that node; for a node joining two individual strains, this is their pairwise correlation coefficient. These values describe profile similarity and are not bootstrap support values. Branches are displayed schematically and are not proportional to clustering distance. Symbol colors show the five k-means group assignments presented in Figure 7, overlaid for comparison. Stars identify the four strains assigned to *Chlorella*—G1093, G1094, G1121, and G1123—and circles identify the remaining strains. H1–H5 identify the five groups obtained by cutting the hierarchy to match the number of PCA-based groups. Agreement between these two partitions of the same physiological dataset was quantified by the adjusted Rand index (ARI = 0.726). The hierarchical group count was fixed for this comparison; see the Supplementary Methods.

### Principal component analysis and k-means clustering

PCA used the same complete matrix of 20 normalized strain-mean profiles presented in Figure 6. Each strain was represented by 34 input variables comprising 17 evolution and 17 respiration values. Evolution and respiration had been normalized separately within each strain, as described above. Variables with variance ≤10⁻¹² were excluded; all 34 variables passed this check. Variables were centered and scaled to unit variance across strains before PCA. Analyses were performed using scikit-learn (Pedregosa et al., 2011). K-means clustering was applied to the unwhitened scores on PC1 and PC2. Candidate solutions with k = 2–10 were evaluated using 10 initializations per cluster count and a fixed random seed of 0. The selected solution maximized the mean silhouette coefficient, calculated using Euclidean distances in PC1–PC2 space (Rousseeuw, 1987). Cluster assignments were displayed on the PCA plot and used to annotate the heatmap for comparison. Individual biological replicates were normalized separately to their own evolution and respiration peaks and projected using the scaler and PCA fitted to the strain means (Figure S8). Replicate points did not determine the PCA axes or the cluster assignments.

The five-group partition was assessed by repeating k-means clustering using the first two, three, four, or five principal components and calculating the adjusted Rand index (ARI) relative to the original two-component partition. Culture-level sensitivity was examined using 2,000 bootstrap resamples, drawing three whole culture profiles with replacement within each strain, recomputing the mean profiles and separate process normalization, and refitting standardization, PCA, and five-group k-means clustering on PC1–PC2 in every resample. Pairwise co-clustering frequencies and each strain’s mean co-clustering frequency with the other members of its original group were calculated without matching numerical cluster labels. Further details are provided in the Supplementary Methods (Figure S9).

Normalized O_2_ evolution–respiration trajectories were constructed using the same strain-level mean profiles analyzed in Figures 6 and 7. Normalized respiration was plotted against normalized evolution, with successive paired observations connected in order of increasing irradiance across the 17 experimental levels. Strains were assigned to panels according to their previously determined k-means cluster membership. For each group, the arithmetic means of normalized evolution and normalized respiration were calculated separately across strains at each corresponding irradiance level, giving each strain equal weight. The resulting paired means were connected in increasing irradiance order to produce the group mean trajectory. All retained strain-level profiles were complete. Individual trajectories were displayed alongside the group means. All panels used identical axis limits and equal scaling of the two axes, retaining negative normalized values. No additional clustering, curve fitting, trajectory simplification, or path-length resampling was performed to construct these plots.

**Figure 7:**
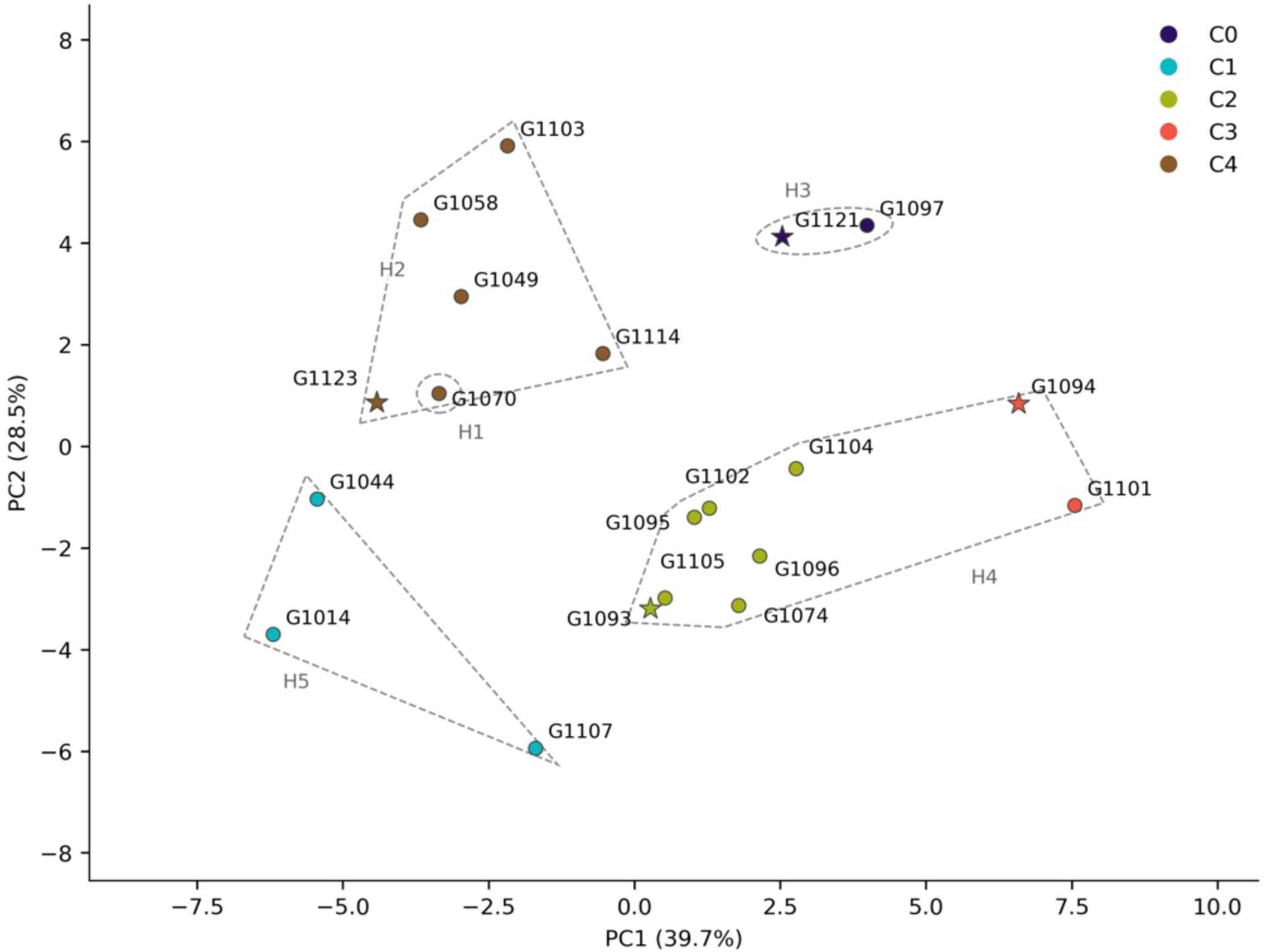
Principal component analysis and k-means clustering of normalized O_2_ evolution and respiration profiles from 20 green microalgal strains. Each point represents one strain, identified by its INCCA-code, based on measurements from three independently grown cultures. Stars identify the four strains assigned to *Chlorella*—G1093, G1094, G1121, and G1123—and circles identify the remaining strains. Mean evolution and respiration profiles were normalized separately within each strain to their respective peak responses, emphasizing relative response patterns. The input comprised the same 34 variables presented in Figure 6, representing the two processes at 17 irradiance levels. Retained variables were centered and scaled to unit variance across strains before PCA. PC1 and PC2 explained 39.7% and 28.5% of the variance, respectively, together accounting for 68.2%. Point colors indicate k-means cluster assignments calculated using PC1–PC2 scores. The clustering analysis identified five groups of strains based on their positions in the PCA plot. Strains positioned closer together have more similar O_2_ exchange response patterns within the two dimensions shown. Different colors distinguish the five groups and match the cluster annotations in Figure 6. Agreement with the five-group hierarchical partition in Figure 6 was ARI = 0.726. Details of clustering, group-count selection, and the ARI calculation are provided in the Supplementary Methods. Grey outlines labelled H1–H5 show the hierarchical memberships from Figure 6 in the PCA projection. They are visual guides to membership, not confidence regions.

### Comparisons in original O_2_ exchange units

For the three pairs sharing named species assignments, the 17 net O_2_ evolution rates and the 17 post-illumination O_2_ consumption rates were plotted separately for each of the three retained cultures per strain in their original concentration-rate units (µmol O_2_ L⁻¹ min⁻¹), without peak normalization. Consumption was expressed as the negative of the recorded dark-interval slope, so positive values indicate O_2_ uptake. Between-strain differences were described by differences between the three-culture means at each irradiance. Complete separation at an irradiance was defined as the lowest rate of one strain exceeding the highest rate of the other. This is a descriptive comparison of the observed culture ranges; culture numbers do not denote matched samples across strains, and irradiance steps are repeated measurements from the same cultures rather than independent replicates.

### Agreement between physiological clustering methods

Agreement between the hierarchical partition in Figure 6 and k-means grouping in Figure 7 was quantified using the adjusted Rand index (ARI; Hubert & Arabie, 1985), implemented in scikit-learn. The average-linkage hierarchy based on the 34-variable correlation distances was cut into exactly five groups to match the selected k-means group count. Assignments were matched by strain identifier before calculating ARI. A value of 1 indicates identical partitions, whereas 0 corresponds to the expected agreement under the chance model used by the index. ARI was reported descriptively without a P value. The five-group hierarchy was used for this comparison and was not independently selected as an optimal partition.

### Association of physiological profiles with phylogeny and taxonomy

Both distance-based analyses used the 20 strain means and the same physiological distance matrix as Figure 6: 1 − Pearson correlation across the 34 concatenated, separately normalized evolution and respiration measurements. No column standardization or PCA reduction was applied to this matrix. Patristic distances were calculated as the sums of branch lengths connecting each pair of study strains in the maximum-likelihood *18S rRNA* tree underlying Figure 5. The 127 reference sequences and the outgroup informed that tree but were not additional physiological observations. A Pearson Mantel test (Mantel, 1967) compared the 190 corresponding off-diagonal distances. Significance was evaluated using 99,999 unrestricted permutations of strain labels, permuting matching rows and columns of the physiological matrix together, with NumPy random seed 0. The two-sided test counted permuted absolute correlations at least as large as the absolute observed correlation.

One-factor PERMANOVA (Anderson, 2001) tested association between physiological distances and genus assignment. *Acutodesmus obliquus* G1102 was assigned to *Tetradesmus* for this analysis to account for its synonymy with *Tetradesmus obliquus*. Pseudo-F and the unadjusted proportion of distance-based sums of squares attributed to genus (R²) were calculated for 12 genera. Significance was evaluated by 99,999 unrestricted permutations of genus labels among strains with random seed 0. Because eight genera were represented by a single strain, the analysis was repeated for the 12 strains belonging to the four genera represented at least twice. Correlation dissimilarities produced negative eigenvalues in principal coordinate analysis; parallel PERMANOVA analyses used sqrt(1 − r), which yielded Euclidean distances. A spatial-median dispersion test (PERMDISP; Anderson, 2006), with small-sample bias adjustment and residual permutations, was performed on these square-root distances for the four replicated genera. This diagnostic does not assess dispersion in singleton genera. Exploratory species-level analyses excluded sp. and cf. assignments, leaving 15 strains in 12 named species and only three residual degrees of freedom.

Permutation P values were calculated as (b + 1)/(99,999 + 1), where b was the number of permuted statistics at least as extreme as the observed statistic. PERMANOVA and PERMDISP used upper-tail tests. P values were unadjusted, and sensitivity analyses were interpreted descriptively. Strains, rather than cultures or pairwise distances, were the units of permutation. The calculations were implemented in Python using NumPy, SciPy, pandas, and Biopython for reading the tree. Full protocols, settings, and supporting results are provided in the Supplementary Methods and Table S5.

## Results

The strains differed in cell size, shape, and colonial organization (Figure 4). Superimposed outlines of individual cells provided a visual representation of typical cell morphology and the variation in cell shape and size within each strain (Figure S1).

The reference-inclusive *18S rRNA* phylogeny placed the study strains among related database sequences (Figures 5 and S7). G1096 and G1107, both identified as *Desmodesmus abundans*, occurred within the same reference-associated clade, supported by 95% of bootstrap replicates. G1044 (*Pediastrum duplex*) and G1070 (*Stauridium tetras*) occupied separate reference-associated branches within a broader clade with 67% support. G1014 (*Chlamydomonas* sp.) and G1103 (*Lobochlamys segnis*) belonged to a clade containing reference sequences with 96% support; a larger clade also containing G1058 (*Haematococcus lacustris*) received 94% support.

All seven study strains assigned to *Chlorella* or *Micractinium* occurred within a broader clade with 100% bootstrap support, although the assigned genera were not resolved as separate groups. The estimated patristic distances among G1093 (*Chlorella ohadii*), G1094 (*Chlorella sorokiniana*), and G1105 (*Micractinium* sp.) were zero in the retained 18S region.

Molecular characterization distinguished morphologically similar strains assigned to different genera. G1074 (*Coelastrella* sp.; Figure 4, panel 2) resembled strains assigned to *Chlorella* and had cell dimensions comparable to those of G1121 (*Chlorella lewinii*; Figure S1). Nevertheless, the reference-inclusive 18S phylogeny placed G1074 in a clade containing G1097, G1102, G1049, G1096, and G1107, supported by 99.2% of bootstrap replicates (Figure 5).

Several separately obtained cultures shared species-level assignments. G1096 and G1107 were both identified as *Desmodesmus abundans*, whereas G1094 and G1123 were both identified as *Chlorella sorokiniana*. G1097 was identified as *Tetradesmus obliquus*, and G1102 was a reference culture supplied as *Acutodesmus obliquus*; these names are taxonomic synonyms referring to the same species (Wynne & Hallan, 2015). The supplied name *Acutodesmus obliquus* was retained for G1102.

Four strains retained genus-level working names: G1014 (*Chlamydomonas* sp.), G1074 (*Coelastrella* sp.), and G1105 and G1114 (*Micractinium* sp.). The species nomenclature of G1104 also remained tentative, as indicated by the designation *Monoraphidium* cf. *dybowskii* (Table 1). To examine how morphological similarity and molecular relationships corresponded to physiological variation, we next compared O_2_ evolution and respiration response profiles across the irradiance series. These comparisons included strains sharing species-level assignments, morphologically similar strains assigned to different genera, and morphologically distinct strains assigned to the same genus.

The retained-reference comparison supported several working identifications but did not independently resolve all species labels (Table S6). In particular, the strongest retained match for G1044, labelled *Pediastrum duplex* from morphology, was *Hydrodictyon reticulatum* OR791100.1 (98.90% identity across the strongest HSP, with 100% query coverage). This discrepancy remains unresolved. The identifications of G1095, G1101, G1104 and G1121 remain working assignments. The retained marker data also did not resolve the generic placement of G1105 and G1114. G1093 was retained as *Chlorella ohadii* on the basis of laboratory culture provenance linking it to SAG 2657; its sequenced rDNA did not distinguish it from *Chlorella sorokiniana* references. Strain codes were therefore retained as the primary identifiers for physiological comparisons.

ITS1 was identical within each nominal species pair: 272 bp for *Chlorella sorokiniana* G1094/G1123, 158 bp for *Desmodesmus abundans* G1096/G1107, and 211 bp for *Tetradesmus obliquus* G1097/G1102. ITS2 was identical over the 204-bp shared region of G1094/G1123 and differed by one substitution in each of the other pairs (231 bp for G1096/G1107; 245 bp for G1097/G1102), with no pairwise indels in these regions. G1123 lacked 25 terminal reference-defined ITS2 positions and G1102 lacked one; these unsequenced positions were excluded from comparisons. No conclusion about compensatory base changes was made (Table S7).

Normalized O_2_ evolution and respiration profiles varied among the 20 strains across the irradiance series (Figure 6). Several pairings in the hierarchical clustering involved strains assigned to different genera. For example, G1104 (*Monoraphidium* cf. *dybowskii*) and G1094 (*Chlorella sorokiniana*) exhibited closely correlated combined profiles (*r* = 0.95), as did G1093 (*Chlorella ohadii*) and G1074 (*Coelastrella* sp.; *r* = 0.94). Conversely, G1096 and G1107, both assigned to *Desmodesmus abundans*, were not recovered as a pair.

To further examine physiological variation among the strains, normalized O_2_ evolution and respiration profiles were analyzed using PCA and k-means clustering (Figure 7). PC1 and PC2 explained 39.7% and 28.5% of the variance, respectively, accounting together for 68.2%. Among the tested cluster counts, the five-cluster solution had the highest mean silhouette coefficient in PC1–PC2 space (0.508). The resulting groups, C0–C4, contained 2, 3, 7, 2, and 6 strains, respectively. G1103 was assigned to C4. The four strains assigned to *Chlorella* were distributed among four different clusters. G1096 and G1107, both assigned to *Desmodesmus abundans*, also belonged to different clusters. These assignments were overlaid on the heatmap to facilitate comparison with the complete normalized response profiles (Figure 6).

The five-group hierarchical partition and PCA-based grouping showed substantial agreement (ARI = 0.726). C0 and C1 were reproduced exactly by hierarchical clustering. The main differences were the combination of C2 and C3 into one hierarchical group and the separation of G1070 from the other five members of C4 (Figures 6 and 7).

The five-group partition was sensitive to the number of retained PCs and to variation among cultures (Figure S9). Relative to the original two-PC partition, ARI was 0.633 using three PCs and 0.510 using either four or five PCs. Across 2,000 within-strain bootstrap resamples, the median ARI was 0.557, with an empirical 2.5th–97.5th percentile range of 0.258–0.954; the complete original partition was recovered in 49 resamples (2.45%). Mean co-clustering with the other members of a strain’s original group ranged from 0.372 to 0.945. The G1094–G1101 pair in C3 co-clustered in 94.45% of resamples, whereas G1107, G1123, and G1114 had mean co-clustering frequencies of 0.372, 0.393, and 0.406 with their original group members, respectively. The five groups therefore summarize patterns in the strain means but are not stable categories for every strain.

The Mantel test comparing 18S patristic distances with physiological distances gave r = 0.103 and P = 0.244 for the 20 strains (99,999 permutations; Table S5).

Genus-level PERMANOVA was not significant (pseudo-F[11,8] = 1.312, R² = 0.643, P = 0.292). Restricting the comparison to 12 strains from four replicated genera gave pseudo-F[3,8] = 0.502, R² = 0.158, and P = 0.728. The square-root-distance analyses were also nonsignificant, as was the corresponding dispersion test for the replicated genera (F[3,8] = 0.453, P = 0.708). Exploratory analyses of the 15 strains with species names lacking sp. or cf. qualifiers likewise detected no clear species effect, but had only three residual degrees of freedom. Full results are reported in Table S5.

Comparison of normalized O_2_ evolution–respiration trajectories revealed variation within and among the five groups identified by PCA and k-means clustering (Figure 8). C0 exhibited relatively high normalized respiration across a broad range of normalized evolution values, although respiration varied among observations with high evolution values. Individual trajectories in C1 and C4 showed changes in direction, indicating that increases in normalized evolution were not consistently accompanied by increases in normalized respiration. C4 also showed considerable variation among strains. C2, the largest group, contained trajectories spanning a broad range of normalized evolution values, with substantial variation in respiration among observations near maximum evolution. In C3, both strains showed an overall increase in normalized respiration alongside increasing normalized evolution, although their individual trajectories differed in curvature. These comparisons describe the observed response patterns without assigning the groups to fitted mathematical curve types.

**Figure 8:**
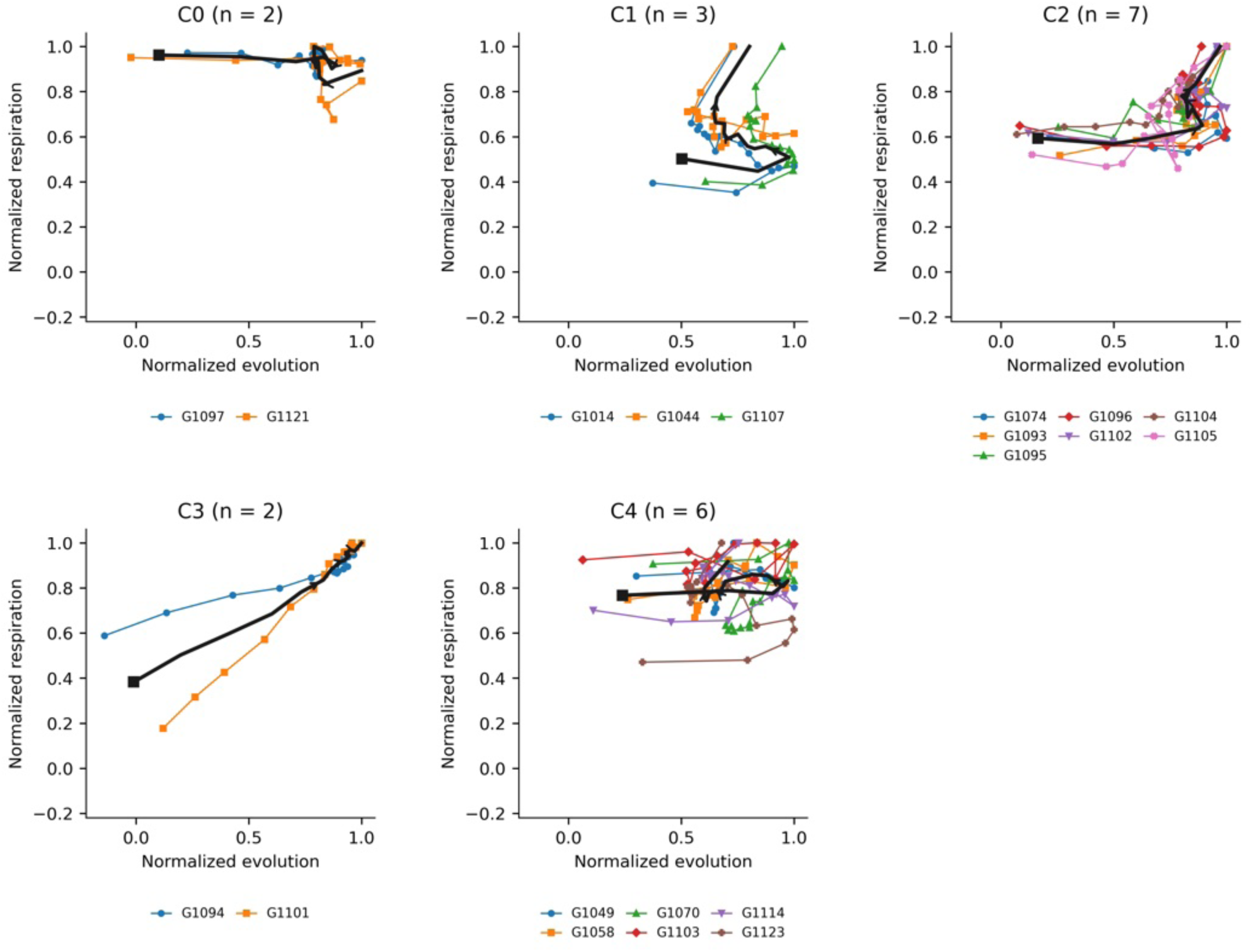
Normalized O_2_ evolution–respiration trajectories of 20 algal strains arranged according to the k-means groups identified in the PCA analysis (Figure 7). Panels C0–C4 contain 2, 3, 7, 2, and 6 strains, respectively. Within each strain, evolution rates were divided by the maximum evolution rate and respiration rates by the most negative respiration rate, setting the strongest response in each process to 1. Negative normalized values indicate responses opposite in sign to the corresponding normalization reference. Normalized respiration is plotted against normalized evolution, with successive observations connected in order of increasing irradiance across 17 levels from 50 to 2000 µmol photons m⁻² s⁻¹. Colors and marker shapes distinguish individual strains, identified by their G-codes. Thick black lines connect the group means calculated at each matching irradiance level, with equal weighting of strains. Black squares mark the group means at the lowest irradiance, and arrowheads indicate progression toward the highest irradiance. All panels use identical axis limits and equal scaling. Panel sample sizes (*n*) indicate numbers of strains. Group membership was inherited from Figure 7; no additional clustering or mathematical curve fitting was performed.

Fixed-window re-extraction covered all 2,040 rates from the 60 retained recordings (Figure S10). Across strains, the median mean absolute change was 1.18% of the original evolution peak and 1.46% of the original respiration peak, but changes were much larger for G1094 (46.9% and 27.5%) and G1101 (43.4% and 41.1%). The fixed-window PCA explained 64.8% of variance in its first two components. At k = 5, agreement with the original grouping was ARI = 0.699, with four strains reassigned. Independent silhouette selection favored four groups (0.452), with ARI = 0.730 relative to the original five-group partition. All three species-assignment pairs remained separated at fixed k = 5, but G1097 and G1102 joined the same group in the selected four-group solution. The direction of the mean difference was preserved at every irradiance for both processes in the *Desmodesmus* and *Tetradesmus* pairs, whereas it was preserved at only 7 of 17 irradiances for evolution and 11 of 17 for respiration in the G1094–G1123 *Chlorella* pair. Thus, rate magnitudes, rankings within this pair, and the detailed grouping depended partly on the extraction window.

The original-unit comparisons showed differences whose magnitude and consistency depended on irradiance (Figure 9). For the *Chlorella sorokiniana* pair, mean net O_2_ evolution at 100 µmol photons m⁻² s⁻¹ was 20.97 µmol O_2_ L⁻¹ min⁻¹ in G1123 and 2.85 in G1094, a difference of 18.12. All three G1123 cultures exceeded all three G1094 cultures at 50, 100, and 150 µmol photons m⁻² s⁻¹, whereas their culture ranges overlapped at the remaining 14 steps. For the *Desmodesmus abundans* pair, G1107 exceeded G1096 at 50 µmol photons m⁻² s⁻¹ (means 9.55 and 1.73, respectively; difference 7.82), with non-overlapping culture ranges. The direction reversed at higher irradiance: all three G1096 cultures exceeded all three G1107 cultures at 300 and at each measured step from 450 to 2,000 µmol photons m⁻² s⁻¹. For the *Tetradesmus*/*Acutodesmus obliquus* pair, all three G1102 cultures exceeded all three G1097 cultures at every measured step from 100 to 2,000 µmol photons m⁻² s⁻¹; the largest mean difference was 10.50 at 300 µmol photons m⁻² s⁻¹ (20.47 versus 9.97). Maximum absolute differences in mean post-illumination O_2_ consumption were 7.37, 3.41, and 8.17 µmol O_2_ L⁻¹ min⁻¹ for the same three pairs, after light exposures of 300, 50, and 2,000 µmol photons m⁻² s⁻¹, respectively. Culture ranges overlapped at the first of these consumption comparisons but were completely separated at the latter two. These descriptions concern the observed cultures and do not constitute tests of statistical significance.

**Figure 9:**
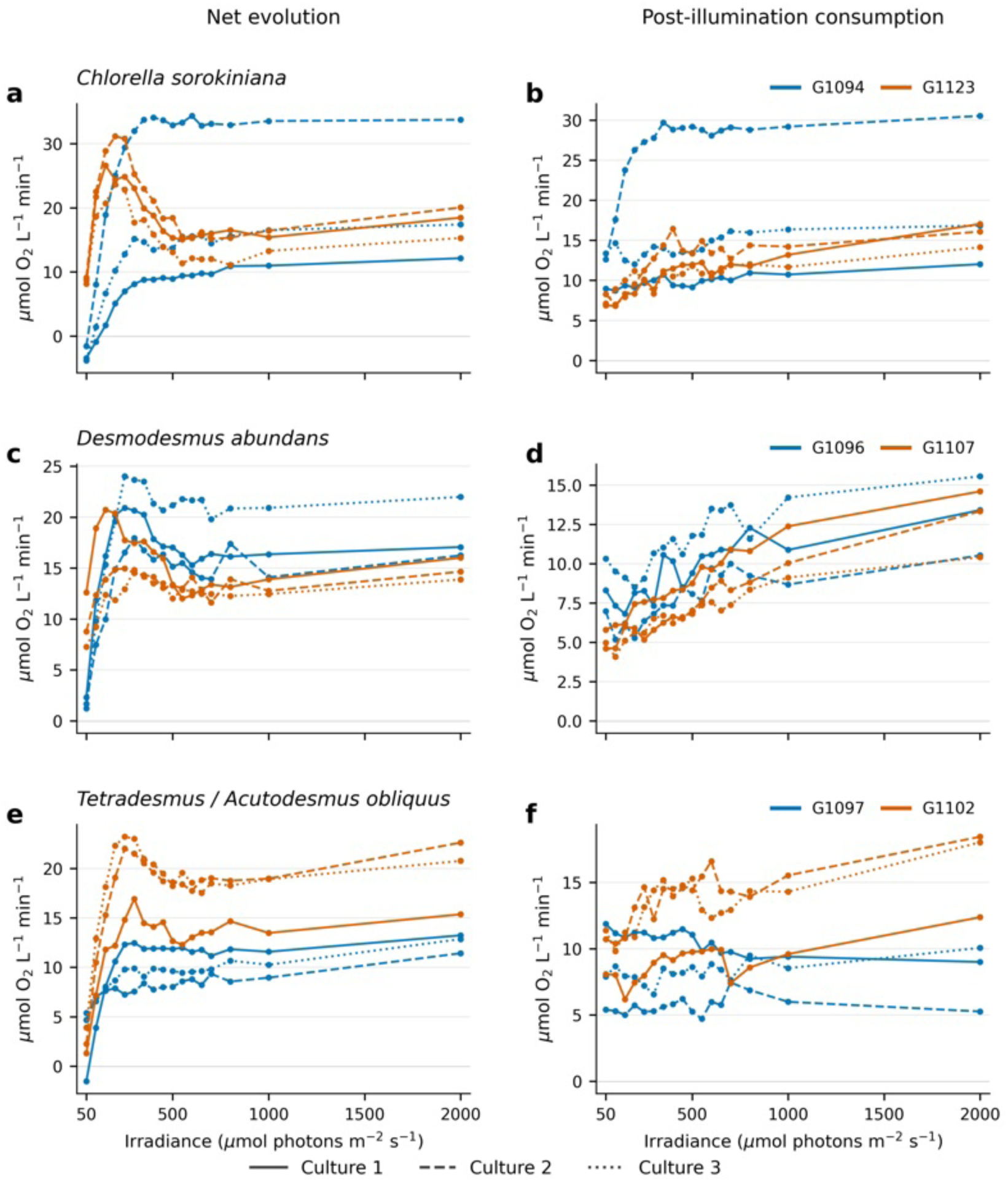
O_2_ exchange in original units for three pairs of strains sharing named species assignments. Net O_2_ evolution (a, c, e) and post-illumination O_2_ consumption (b, d, f) are plotted against irradiance for *Chlorella sorokiniana* (G1094, blue; G1123, orange; a, b), *Desmodesmus abundans* (G1096, blue; G1107, orange; c, d), and *Tetradesmus*/*Acutodesmus obliquus* (G1097, blue; G1102, orange; e, f). Each line connects the 17 measured rates from one independently grown culture; solid, dashed, and dotted lines distinguish the three cultures within a strain and do not indicate pairing between strains. Cultures were measured at a common loading of 20 µg total chlorophyll mL⁻¹. Rates are expressed in µmol O_2_ L⁻¹ min⁻¹ without peak normalization; consumption is the negative of the recorded dark-interval slope, so positive values indicate O_2_ uptake. Each consumption rate is plotted at the irradiance of the immediately preceding light interval. Connecting lines show the experimental order and are not fitted response curves. Negative net evolution values are retained.

## Discussion

This study combined morphological observations, rDNA-based identification, and O_2_ exchange measurements to characterize 20 green microalgal strains. Microscopy documented cellular and colonial features, molecular analysis provided taxonomic context, and physiological measurements revealed differences in relative responses to increasing irradiance. The three pairs sharing named species assignments occupied different groups in the analysis of strain means. The genus-level analysis detected no clear association between assigned genus and physiological distances.

Morphological characterization provided an initial framework for distinguishing the strains. Micrographs and superimposed cell outlines revealed differences in cell shape, size, and colonial organization (Figure 4; Figure S1). Nevertheless, similar morphology occurred among strains assigned to different genera. G1074, identified as *Coelastrella sp.*, resembled strains assigned to *Chlorella* despite their placement in different classes of *Chlorophyta*. Its association with Scenedesmaceae strains in Figure 5 is consistent with the classification of *Coelastrella* (Goecke et al., 2020).

The assignment of G1101 (*Chlorolobion braunii*) to the Selenastraceae family is consistent with chloroplast-genome evidence for *C. braunii* (Mu et al., 2023). The reference-associated clade containing G1014 (*Chlamydomonas* sp.) and G1103 (*Lobochlamys segnis*) reflects the historically broad circumscription of *Chlamydomonas*, which led to the establishment of *Lobochlamys* (Pröschold et al., 2001).

Adding database reference sequences improved the taxonomic context of the phylogeny, but several relationships remained poorly resolved. The strains assigned to *Chlorella* and *Micractinium* were not separated into distinct generic groups, and several study strains had zero estimated patristic distances in the retained 18S region. Such distances indicate that the fitted tree did not resolve divergence between those strains, not that they share a species. Bootstrap support reflects consistency under resampling of alignment positions, not the probability that an identification is correct (Felsenstein, 1985). Morphological observations, culture provenance and ribosomal markers provided complementary identification evidence, but neither a high percentage identity in a conserved HSP nor placement in a broad 18S clade alone established species identity. The unresolved morphological/molecular discrepancy for G1044 and the provisional names of several other cultures limit species-level and genus-level interpretations. The close within-pair ITS similarities are consistent with the paired working assignments, but they do not demonstrate reproductive compatibility or exclude cryptic diversity.

These uncertainties also clarify the role of physiological characterization. Shared species assignments do not establish that separately obtained strains are genetically identical or physiologically interchangeable. The O_2_ exchange assay addressed a complementary question: how did the strains respond to the same experimental irradiance sequence? Standardizing chlorophyll loading provided a common basis for these measurements, while normalization emphasized the relative changes in each response profile (Figure 6). Normalized profiles describe response patterns relative to each strain’s own peak, whereas the original-unit comparisons retain rate magnitude and biological replicate variation (Figure 9).

The physiological analyses revealed variation that was not consistently captured by the assigned taxonomic groups (Figures 6 and 7). G1096 and G1107, both assigned to *Desmodesmus abundans*, occupied different PCA clusters. The same was true for G1094 and G1123, both assigned to *Chlorella sorokiniana*. G1097 and G1102 likewise occupied different clusters, although the two names under which they were obtained, *Tetradesmus obliquus* and *Acutodesmus obliquus*, are synonyms referring to the same species (Wynne & Hallan, 2015). Thus, all three pairs sharing the same species nomenclature were placed in different groups based on their mean physiological profiles; the two cultures labelled *Micractinium* sp. were not treated as confirmed members of the same species. Furthermore, the four strains assigned to *Chlorella* were distributed among four of the five clusters.

Several explanations remain compatible with the physiological differences among strains sharing species assignments. They may reflect intraspecific functional variation, consistent with the substantial differences in genome content reported among strains carrying the name *Chlorella sorokiniana* (Hovde et al., 2018). Alternatively, some assignments may conceal distinct lineages that the available ribosomal data do not resolve. Species boundaries within the *Chlorella* clade have been revised as molecular evidence accumulated (Bock et al., 2011; Krivina & Temraleeva, 2020), and comparable identification difficulties have long been recognized in *Scenedesmus* and related genera (Hegewald & Silva, 1988; Trainor & Egan, 1991). ITS1 was identical within each pair, while ITS2 differed by zero or one substitution over the available shared regions (Table S7). These similarities are compatible with the working assignments but do not resolve species boundaries; partial ITS2 coverage in two cultures and the absence of a secondary-structure comparison further limit interpretation (Darienko et al., 2015).

PCA and k-means provided an exploratory summary of this physiological variation. The first two principal components accounted for 68.2% of the variance in the standardized normalized profiles, and the selected five-cluster solution had a mean silhouette coefficient of 0.508 (Figure 7). However, the original partition was recovered exactly in only 49 of 2,000 within-strain bootstrap resamples (2.45%), and agreement declined when additional principal components were used (Figure S9). These results support using C0–C4 as descriptive labels for the displayed mean profiles, rather than as reproducible physiological classes.

Projection of the 60 individual biological replicates into the same PCA space revealed substantial differences in dispersion among strains (Figure S8). Replicates from some strains occupied relatively compact regions, whereas others were widely spread. Each culture was normalized to its own evolution and respiration peaks, so the dispersion reflects variation in relative profile shape, although normalization can accentuate variation when a peak rate is small. Separation between two strain means therefore does not necessarily imply consistently distinct responses among individual cultures. The isolated G1105 replicate (10 February 2026, repeat 1) had a respiration normalization denominator of −2.68 µmol O_2_ L⁻¹ min⁻¹, compared with −12.25 and −10.54 in the other two cultures. Positive slopes in 13 of its 17 nominal dark intervals produced negative normalized respiration values, which were amplified by the small denominator (Figure S8). This replicate was retained; the recording alone does not establish whether its atypical profile arose from biological variation or measurement conditions.

The ARI comparison quantified the extent to which two analyses of the same physiological measurements produced similar groupings. Agreement was substantial at a matched count of five groups (ARI = 0.726). The ARI does not test agreement with taxonomy or phylogeny and does not measure reproducibility across biological replicates.

Comparison of the physiological groupings with cell dimensions revealed partial correspondence in individual cases, although the groups did not form uniform size classes (Figure S1; Table 2). Two cultures assigned to *Micractinium*, G1095 and G1105, had similar mean dimensions of 4.73 × 3.76 and 4.60 × 3.85 µm, respectively, and both belonged to C2 and H4. These groups also included the small-celled *Chlorella ohadii* G1093, providing a local overlap between small cell size, related taxonomic assignments within Chlorellaceae, and physiological similarity. However, the similarly small *Micractinium* G1114 belonged to C4 and H2. Those groups also contained strains with markedly different dimensions, including *Haematococcus lacustris* G1058 (27.54 × 26.61 µm) and *Chlorella sorokiniana* G1123 (4.55 × 4.04 µm). Conversely, G1097 and G1102, supplied under synonymous names for *Tetradesmus obliquus*, had similar mean dimensions but occupied different C and H groups.

**Table 2:** assignment of each algal strain INCCA code to its C group according to figure 7 and H group according to figure 6.

| **Strain INCCA Code** | **C group (Figure 7)** | **H Group (Figure 6)** |
|---|---|---|
| G1014 | C1 | H5 |
| G1044 | C1 | H5 |
| G1049 | C4 | H2 |
| G1058 | C4 | H2 |
| G1070 | C4 | H1 |
| G1074 | C2 | H4 |
| G1093 | C2 | H4 |
| G1094 | C3 | H4 |
| G1095 | C2 | H4 |
| G1096 | C2 | H4 |
| G1097 | C0 | H3 |
| G1101 | C3 | H4 |
| G1102 | C2 | H4 |
| G1103 | C4 | H2 |
| G1104 | C2 | H4 |
| G1105 | C2 | H4 |
| G1107 | C1 | H5 |
| G1114 | C4 | H2 |
| G1121 | C0 | H3 |
| G1123 | C4 | H2 |

Thus, neither similar cell size nor a shared taxonomic assignment consistently predicted physiological group membership. The C and H partitions summarize similarities in normalized O₂ exchange profiles, while the morphometric observations came from one culture per strain. Their correspondence should therefore be interpreted descriptively and does not establish a causal relationship between cell size and physiological response.

The Mantel test provided little evidence of an association between 18S patristic distances and differences in normalized O₂ exchange profiles. The comparison reflects the resolution of the retained 18S positions and the physiological responses measured in this collection.

The PERMANOVA results provided no clear evidence that physiological profiles were grouped by assigned genus. The all-strain model attributed 64.3% of the distance-based sums of squares to genus, but that value should not be interpreted as strong predictive performance: 12 genus groups were fitted to only 20 strains, and eight groups contained a single strain. Among the four genera represented by multiple strains, the corresponding R² was 0.158 and P = 0.728.

The assigned genera did not clearly organize the measured physiological differences in this collection. The data are most informative as evidence that the measured strain means contain functional information that is not fully conveyed by the assigned names.

The evolution–respiration trajectories made the physiological variation easier to interpret by displaying paired responses in their experimental order (Figure 8). For example, C0 maintained relatively high normalized respiration across a broad range of normalized evolution values, whereas C1 and the six-member C4 contained trajectories with pronounced changes in direction. C3 showed an overall increase in both normalized responses, although the individual trajectories differed in curvature. Respiration values near 1 indicate proximity to a strain’s own maximum response, rather than greater absolute respiration than in another group. Because an average can conceal variation, the individual trajectories remain essential; a common cluster label does not imply a uniform response.

Interpretation of these trajectories must also account for the duration and sequence of the measurements. Respiration measured immediately after illumination may depend on the preceding light exposure. Light-enhanced dark respiration has been demonstrated in *Chlamydomonas reinhardtii*, with post illumination O_2_ consumption varying with irradiance and illumination duration (Xue et al., 1996). This provides a possible physiological context for changes in the respiration profiles, without establishing that the same mechanism explains the differences among the present strains. Inorganic carbon availability can also alter O_2_ exchange: in *Chlorella sorokiniana*, carbon depletion reduced net O_2_ evolution, whereas bicarbonate addition restored it (Patil et al., 2020). Carbon depletion was not measured here, so its contribution to the present response profiles remains unresolved. The assay used TAP without added bicarbonate, and neither pH nor dissolved inorganic carbon was measured. The fixed-window sensitivity analysis assesses dependence on rate extraction but cannot separate irradiance effects from time, O_2_ accumulation, or carbon availability.

Culture provenance showed partial correspondence with the physiological groupings. Four of the six strains in C4 were obtained from INCCA, whereas C0 and C3 contained only strains isolated directly in our laboratory. However, INCCA strains also occurred in C1 and C2, and C4 included strains with reported isolation dates ranging from 1974 to 2026. Also, from various geographical locations with different biomes. Thus, C4 did not represent a uniform group of long-maintained cultures.

A species assignment provides useful biological context, but the measurements here show why individual strains should be characterized when their physiological responses matter. The original-unit comparisons reveal both differences in rate magnitude and variation among independently grown cultures (Figure 9). The sensitivity of some comparisons and group assignments to the extraction window, together with limited molecular resolution, constrains their generalization. The present profiles describe responses under a common assay sequence; they do not establish steady-state capacities, productivity rankings, or suitability for a particular application.

Limitations. (i) The collection comprises 20 strains, only three pairs of which share a named species assignment. The assignments rely on morphology and rDNA similarity, and neither 18S proximity nor the available ITS comparisons establish conspecificity. (ii) Each strain was measured in three independently grown cultures during one ascending irradiance series. Irradiance is confounded with elapsed time, accumulated O_2_, and possible carbon depletion; pH, dissolved inorganic carbon, and cell counts were not measured. The cultures were grown at 2% CO_2_, then assayed in TAP without supplied CO_2_ or added bicarbonate. The short selected slopes are operational response measurements, not steady-state capacities. (iii) Separate peak normalization removes absolute magnitudes and the respiration-to-evolution ratio, while equal chlorophyll loading does not imply equal cell number or biomass composition. (iv) Morphology was recorded from one culture per strain, and culture history cannot be separated from strain identity. (v) Axenicity was not tested, so bacterial O_2_ consumption cannot be excluded. (vi) Resampling, additional principal components, and fixed-window extraction changed some group assignments; the five groups remain descriptive. No carbon-supplementation or reversed/randomized-light-order control was performed, so these confounding effects remain unresolved.

## Conclusions

Combining microscopy, rDNA analysis, and O_2_ exchange measurements provided complementary information about 20 strains from the phylum Chlorophyta. Strains sharing named species assignments differed in their measured response profiles and original-unit rates, but variation among cultures and sensitivity to rate extraction limit generalization. The five PCA-based groups summarize the original strain means and are not established physiological classes. Physiological distances showed no clear association with 18S patristic distances or assigned genus. The study supports direct strain-level physiological characterization alongside taxonomic identification under clearly stated assay conditions; it does not establish confirmed intraspecific differentiation or independence of physiology from evolutionary history.

## Data availability

The rDNA contigs, retained raw O_2_ recordings, extracted rates, sequence alignments, phylogenetic trees, and analysis code are provided in the accompanying data and code package. Available study-sequence GenBank accession records are listed in Table S6. The remaining pending accession and the permanent public repository identifier must be added after deposition. Requests for study cultures should be directed first to Ruth N. Kaplan-Levy, Head of INCCA, subject to culture availability and applicable transfer arrangements.

## Acknowledgments

We thank the Yacoby laboratory members for assistance during this study, and Eyal Zaksh and Eden Zloof for obtaining algal samples.

## Author contributions

M.K. and I.Y. conceived the study, designed the experiments, analyzed the data, and wrote the manuscript. I.Y. and R.N.K.-L. supervised the work. I.Y. secured funding. T.S., S.M., D.P., L.O.-Y., and R.N.K.-L. contributed to the research. A.A isolated the strains originated from INCCA.

AI assistance. OpenAI Codex was used for language editing, development of analysis scripts, and preparation of document files. The analysis scripts and numerical outputs are included in the accompanying data and code package.

