## Supplemental for "O_2_ exchange profiling reveals physiological diversity among taxonomically characterized green microalgal strains"

| Index | INCCA Code | Identification | AlgaeBase ID | Sample Region | Sample Location | Latitude | Longitude | Sample year |
|---|---|---|---|---|---|---|---|---|
| **1** | G1014 | *Chlamydomonas* sp. | 43319 | Lake Kinneret | Station A | 32.821299 | 35.589042 | 2012 |
| **2** | G1044 | *Pediastrum duplex* | 27511 | Lake Kinneret | Station A | 32.821299 | 35.589042 | 2014 |
| **3** | G1049 | *Desmodesmus denticulatus* | 42405 | Lake Kinneret | Station A | 32.821299 | 35.589042 | 2018 |
| **4** | G1058 | *Haematococcus lacustris* | 36266 | Lake Kinneret | Station A | 32.821299 | 35.589042 | 1974 |
| **5** | G1070 | *Stauridium tetras* | 129243 | Lake Kinneret | Station A | 32.821299 | 35.589042 | 2020 |
| **6** | G1074 | *Coelastrella* sp. | 45299 | Israel National Water Carrier | Eshkol Reservoir | 32.77549 | 35.26004 | 2000 |
| **7** | G1093 | *Chlorella ohadii* | 177095 | North-West Negev | Nitzana Dunes (Biological sand crust) | 30.933333 | 34.383333 | 2012 |
| **8** | G1094 | *Chlorella sorokiniana* | 51508 | Tel Aviv-Yafo | Yarkon River | 32.09623173 | 34.78216334 | 2022 |
| **9** | G1095 | *Micractinium tetrahymenae* | 176401 | Tel Aviv-Yafo | Yarkon River | 32.09623173 | 34.78216334 | 2022 |
| **10** | G1096 | *Desmodesmus abundans* | 42384 | Lake Kinneret | Kursi Beach | 34.492929 | 35.384136 | 2023 |
| **11** | G1097 | *Tetradesmus obliquus* | 158061 | Negev | Ein Yorqe'am | 31.03166667 | 35.04138889 | 2024 |
| **12** | G1101 | *Chlorolobion braunii* | 27677 | Negev | Ein Yorqe'am | 31.03166667 | 35.04138889 | 2024 |
| **13** | G1102 | *Acutodesmus obliquus* SAG 276-3d | 135513 | Italy | Castellamare | 40.706018 | 14.483042 | 1954 |
| **14** | G1103 | *Lobochlamys segnis* | 118596 | Golan Heights | El-Al Stream | 32.82861111 | 35.89888889 | 2023 |
| **15** | G1104 | *Monoraphidium* cf. *dybowskii* | 131950 | Yarkon River | Outflow | 32.09623173 | 34.78216334 | 2022 |
| **16** | G1105 | *Micractinium* sp. | 43421 | HaSharon | Hadera Stream Park | 32.46805556 | 34.90305556 | 2023 |
| **17** | G1107 | *Desmodesmus abundans* | 42384 | Golan Heights | Winter puddle near Bental reservoir | 33.23722222 | 36.30527778 | 2023 |
| **18** | G1114 | *Micractinium* sp. | 43421 | Arava | Yotveta Reservoir | 29.8847343 | 35.06469748 | 2024 |
| **19** | G1121 | *Chlorella lewinii* | 147234 | Modi’in | Modi’in | 31.535084 | 34.590826 | 2025 |
| **20** | G1123 | *Chlorella sorokiniana* | 51508 | Negev | Gev Yamin | 30.943929 | 35.073108 | 2026 |

**Table S1:** Working identifications and recorded isolation locations. Strains 1–6 and 18 are from INCCA; 8–12, 14–17 and 19–20 were isolated in the Yacoby laboratory. G1102 was supplied as *Acutodesmus obliquus* SAG 276-3d. Laboratory provenance links G1093 to *Chlorella ohadii* SAG 2657; it was not obtained directly from SAG.

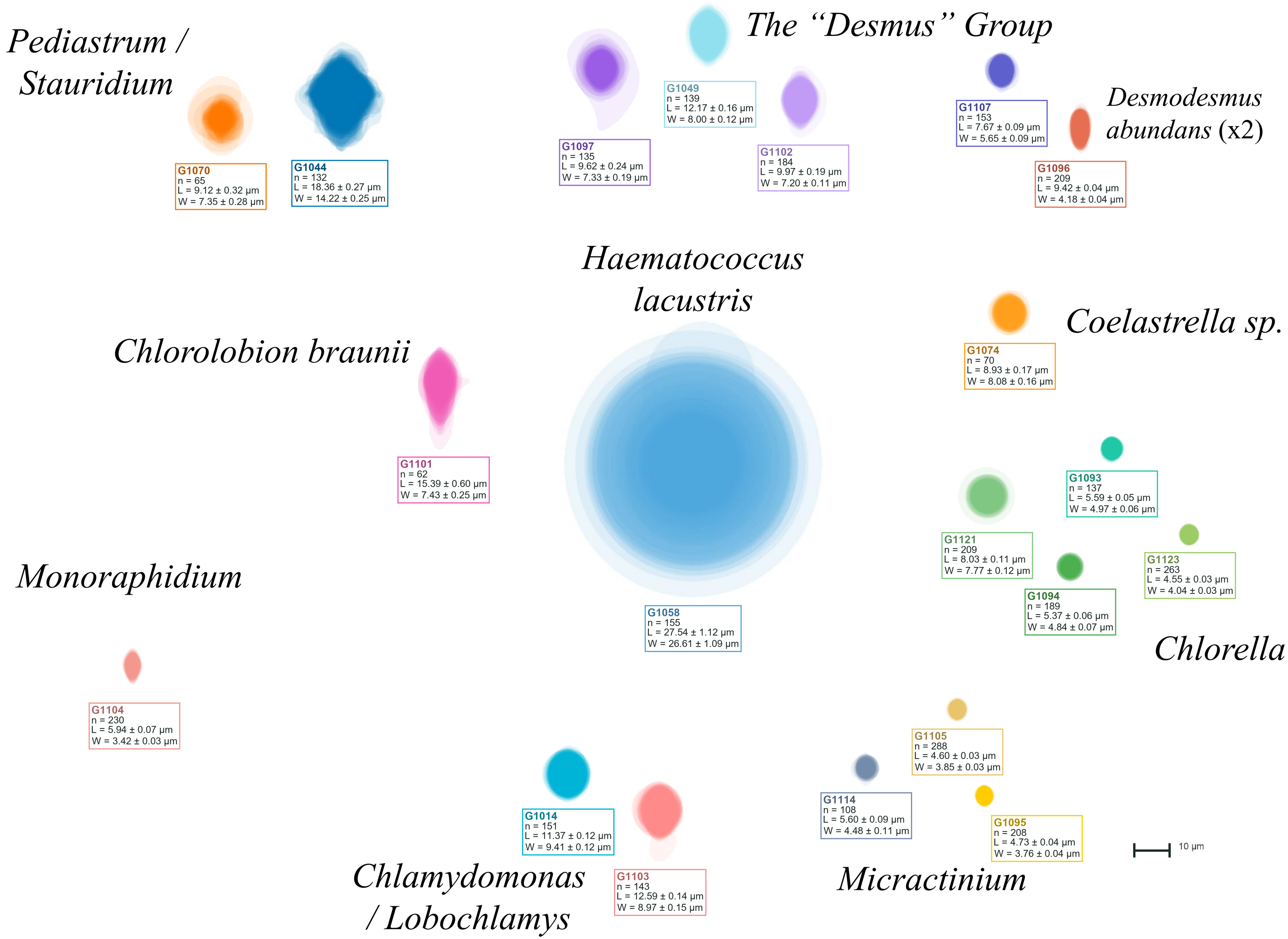


**Figure S1:** Superimposed cell outlines for the 20 green microalgal strains measured using Algal Detector. Each label shows the number of measured cell images (n) and mean length (L) and width (W) ± standard error. Cells were measured from one culture per strain. G1123 is shown in the *Chlorella* group. Scale bar, 10 µm.

| No. | Name | Sequence 5' → 3' |
|---|---|---|
| **1** | 18S-ITS 1st - NS1 | GTA GTC ATA TGC TTG TCT C |
| **2** | 18S-ITS 1st - ITS4 | TTC CTC CGC TTA TTG ATA TGC |
| **3** | 18S rDNA 2nd - 18Sseq | GCT CGT AGT TGG ATT TCG GGT GGG |
| **4** | 18S rDNA 2nd - ITSseq | CAC TAC GTA TCG CAT TTC GCT G |

**Table S2:** Primers used for amplification and Sanger sequencing of the nuclear ribosomal region spanning 18S, ITS1, 5.8S, ITS2, and part of 28S. Primers 1 and 2 were used for PCR amplification; primers 1–4 were used for Sanger sequencing.

| No. | Name | Start | Cycles Denaturation | Annealing | Extension | End | Cycles | Primer source |
|---|---|---|---|---|---|---|---|---|
| **1** | 18S-ITS 1st – NS1 | 95°C, 2–3 min | 95°C, 30 sec | 59°C, 30 sec | 72°C, 0.5–2 min | 72°C, 10 min | 36 | White et al. (1990) |
| **2** | 18S-ITS 1st – ITS4 | | | | | | | |
| **3** | 18S rDNA 2nd – 18Sseq | Used For Sanger Sequencing | | | | | | Kaplan-Levy et al. (2016) |
| **4** | 18S rDNA 2nd – ITSseq | | | | | | | |

**Table S3:** PCR conditions for primer pair 1/2 shown in Table S2. Each reaction contained VeriFi Hot Start Mix (PCR Biosystems; 12.5 µL in a 25-µL reaction or 25 µL in a 50-µL reaction), 0.2 µM primers, and 50–100 ng template DNA. Primers 3 and 4 were used for Sanger sequencing rather than PCR amplification.

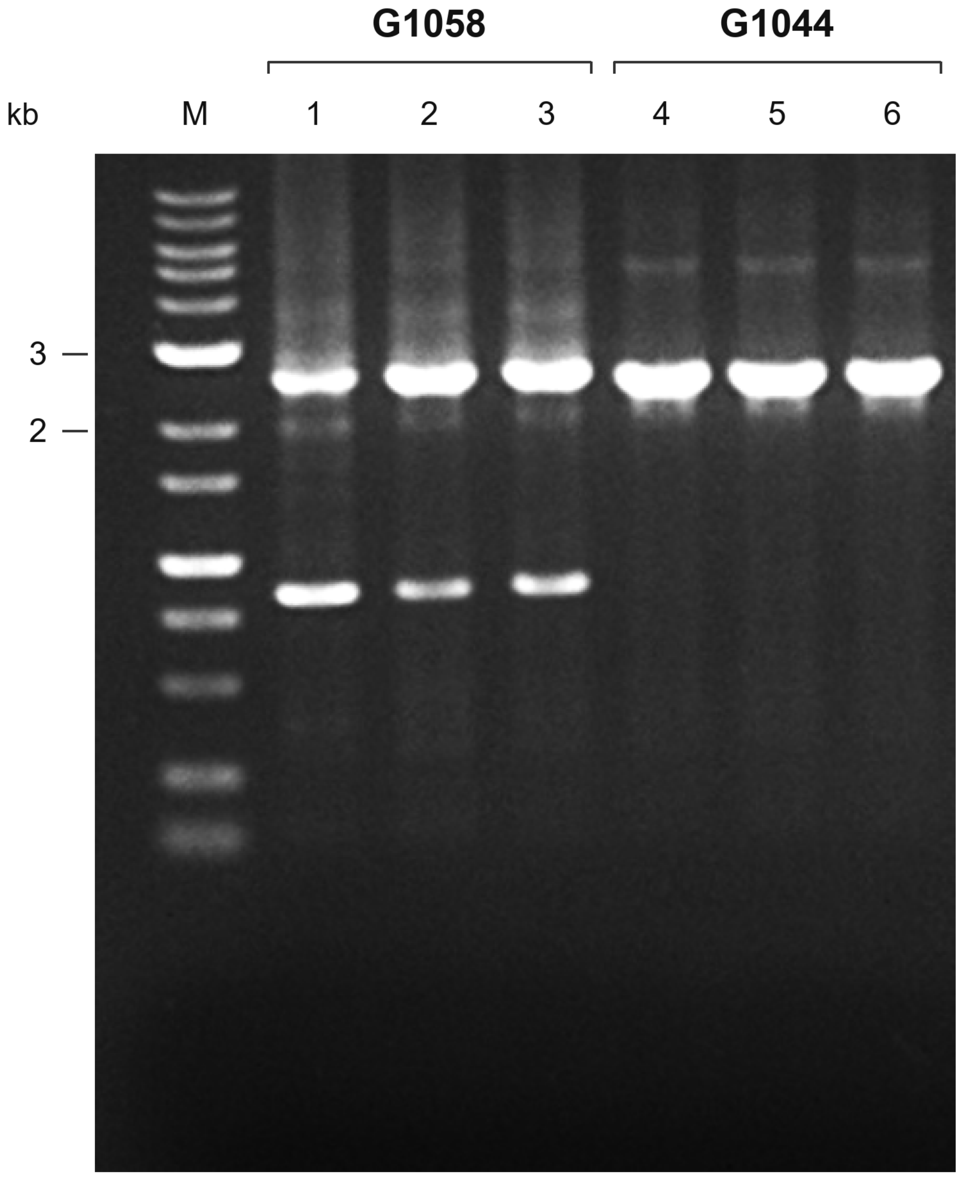


**Figure S2:** Agarose gel electrophoresis of ribosomal DNA amplification products from G1058 and G1044. PCR products generated using primers 1 and 2 were separated on a 1% (w/v) agarose gel. Lanes 1–3, G1058; lanes 4–6, G1044; M, 1-kb DNA size marker. Prominent upper bands migrated between the indicated 2 and 3 kb markers. Because G1058 yielded multiple bands, the selected band was excised and purified from the agarose gel before Sanger sequencing. The image is a continuous crop showing the marker and six adjacent sample lanes.

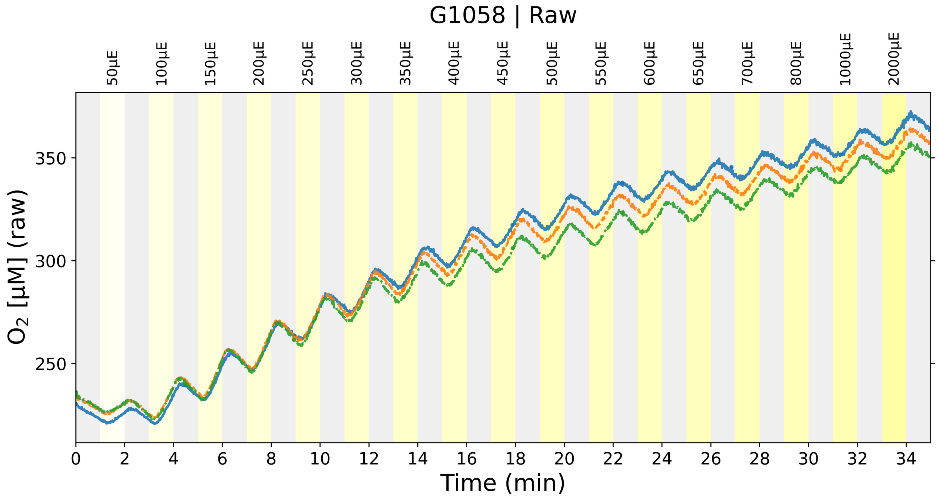


**Figure S3:** FireSting result for microalgal strain INCCA-G1058. Dissolved $O_2$ concentration (µM) was recorded over time during alternating 1min dark and light intervals. Light intervals were applied at stepwise increasing irradiance levels ranging from 50 to 2000 µmol photons $m^{-2}$ $s^{-1}$. Grey and yellow shaded regions represent dark and light periods, respectively. Three biological replicates are shown, each line represent one replicate.

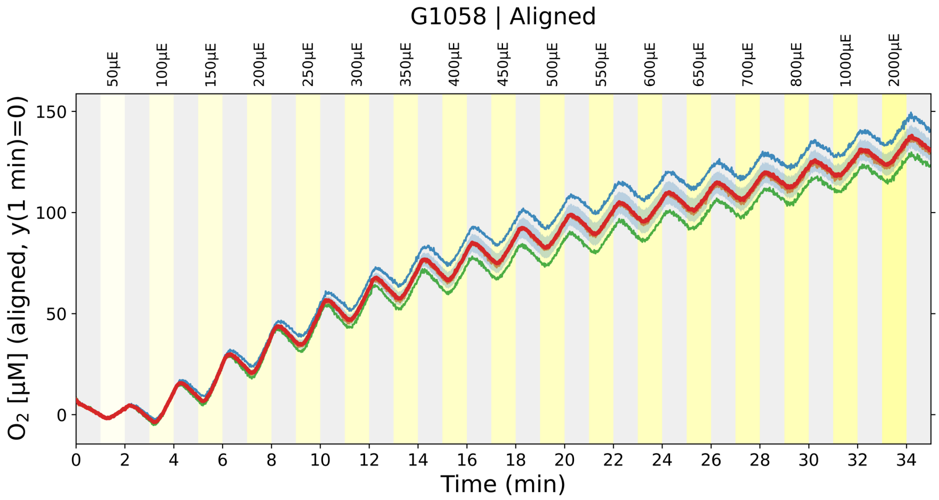


**Figure S4:** Aligned FireSting result for microalgal strain INCCA-G1058. Dissolved $O_2$ concentration (µM) was recorded over time during alternating 1 min dark and light intervals under stepwise increasing irradiance (50-2000 µmol photons $m^{-2}$ $s^{-1}$). Each trace was aligned by setting the $O_2$ concentration at 1 min to zero. Grey and yellow shaded regions represent dark and light periods, respectively. The red curve represents the average of three biological replicates (blue, green and orange).

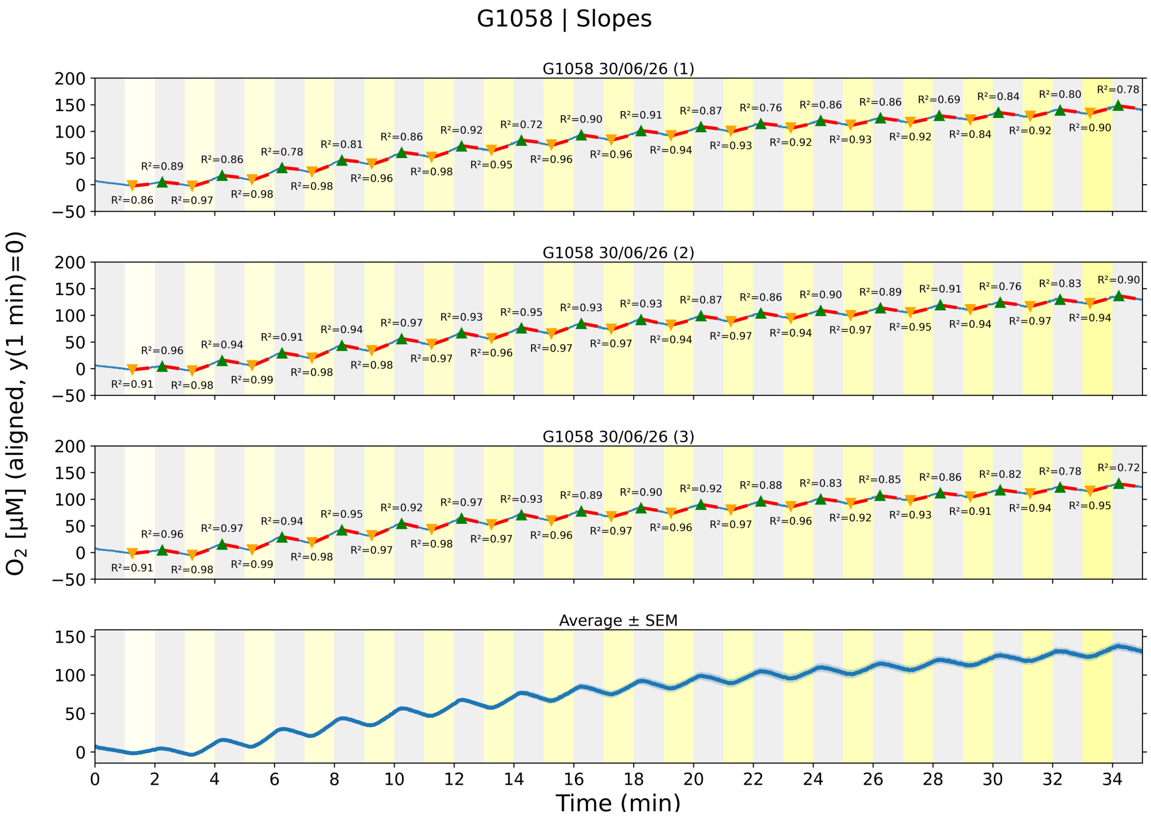


**Figure S5:** Evolution and respiration slopes for microalgal strain INCCA-G1058. Aligned $O_2$ traces (y(1 min) = 0) from three biological replicates are shown. Rates were fitted over 30 s windows whose starts ranged from 15 s before to 15 s after each nominal light or dark transition, in 1.2 s increments. The maximum slope was selected for light-associated $O_2$ evolution and the minimum slope for dark-associated $O_2$ consumption; candidate windows could cross a nominal transition. The bottom panel shows the mean trace with its standard error. Irradiance increased sequentially from 50 to 2000 µmol photons $m^{-2}$ $s^{-1}$. A separate analysis using the same fixed central 15–45 s window in every interval is presented in Figure S10.

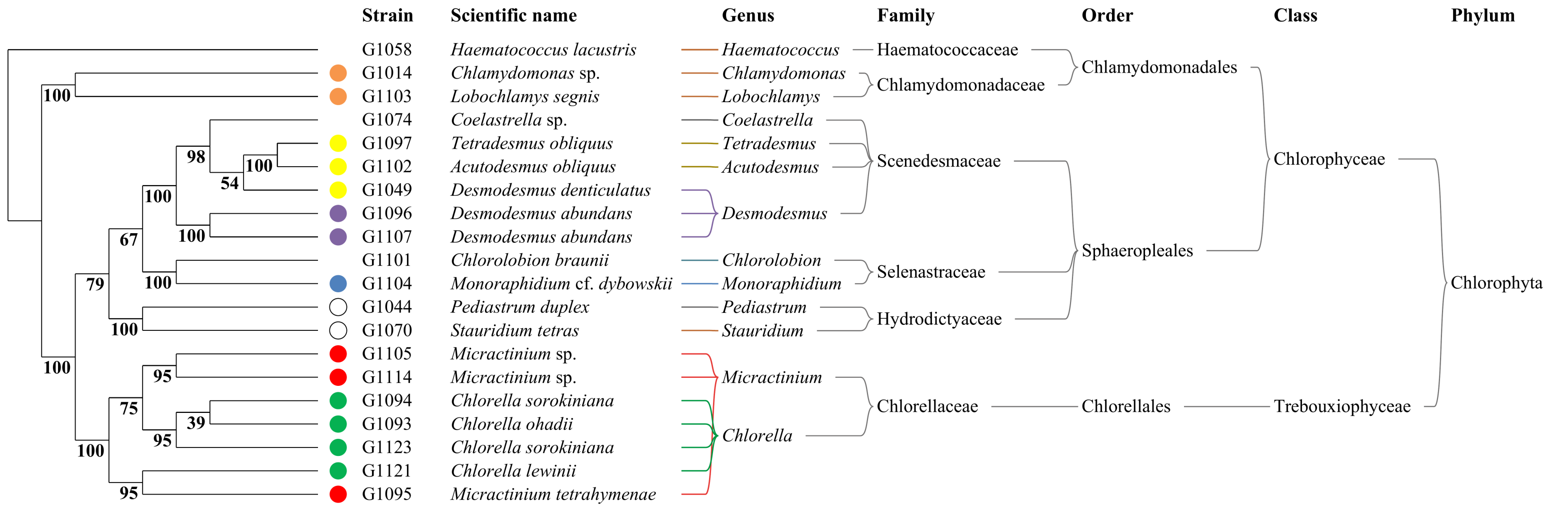


**Figure S6:** Maximum-likelihood phylogeny and taxonomic classification of the 20 green microalgal strains used in this study, including the previously identified model strains G1093 and G1102. The maximum-likelihood tree inferred in MEGA12 from aligned ribosomal DNA sequences is shown (log likelihood −8,724.24). Tip labels identify strains by G-code, with their assigned scientific names shown alongside; "sp." indicates identification to the genus level, and "cf." indicates a tentative species-level assignment. Bold numbers at internal nodes indicate bootstrap support (%) from 1,000 replicates. Branches are displayed schematically and are not proportional to evolutionary distance. No outgroup was specified; the displayed root does not establish evolutionary direction. Colored circles indicate the predefined groups used in Figure S1: yellow, *Tetradesmus obliquus*, *Acutodesmus obliquus*, and *Desmodesmus denticulatus*; purple, *Desmodesmus abundans*; blue, *Monoraphidium*; white, *Pediastrum/Stauridium*; red, *Micractinium*; green, *Chlorella*; and orange, *Chlamydomonas/Lobochlamys*. G1074, G1101, and G1058, which are presented individually in Figure S1, have no colored circles. The right-hand diagram links the assigned scientific names to their genera through colored connectors and summarizes their classification by family, order, class, and phylum. It represents taxonomic classification rather than a separately inferred phylogeny; connector lengths and crossings have no evolutionary meaning. The name *Acutodesmus obliquus* is retained for G1102, and is also shown in the genus column. This supplied name is a synonym of *Tetradesmus obliquus*, the name used for G1097.

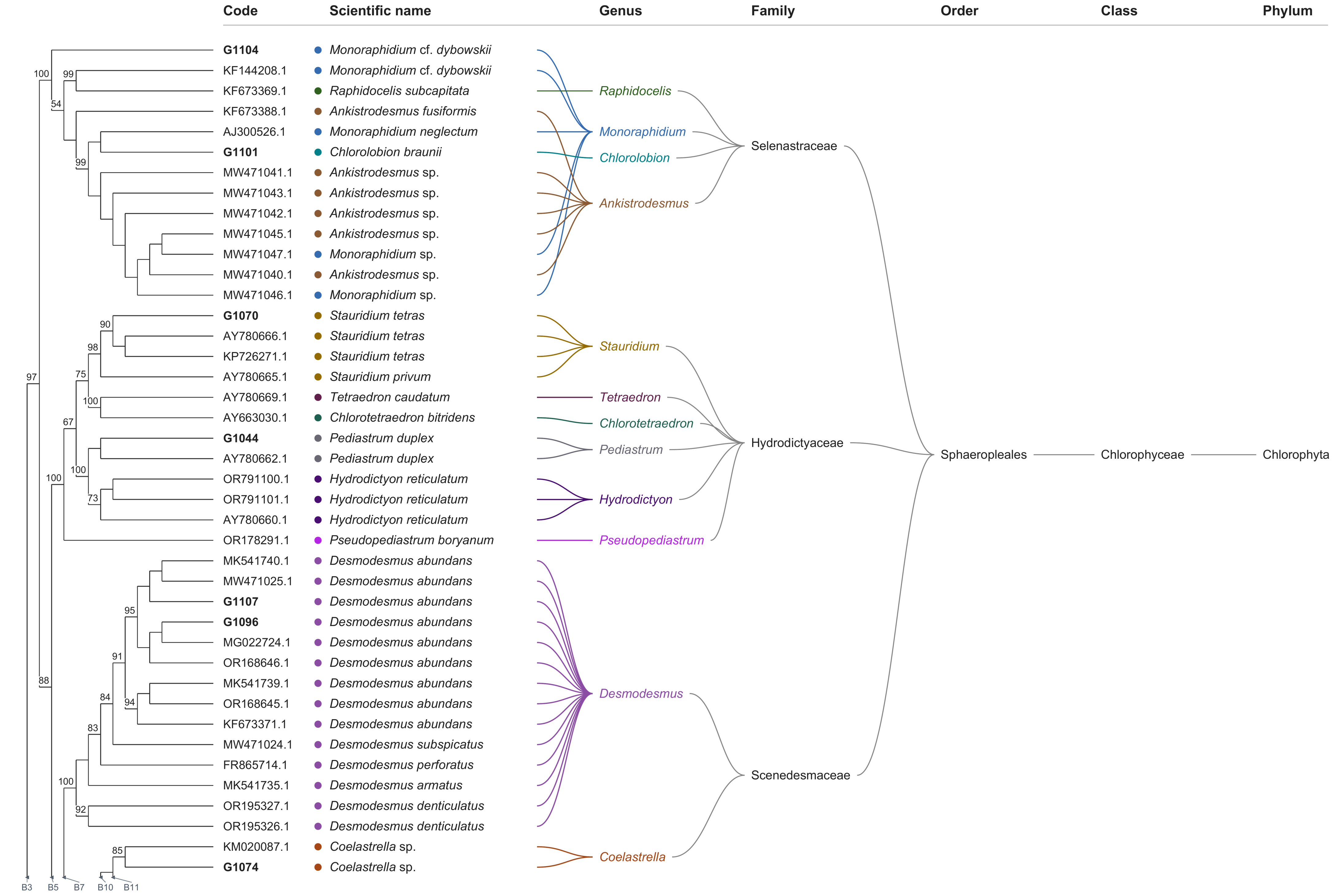


Complete tree - part 1 of 4

Tips 1-41 of 148

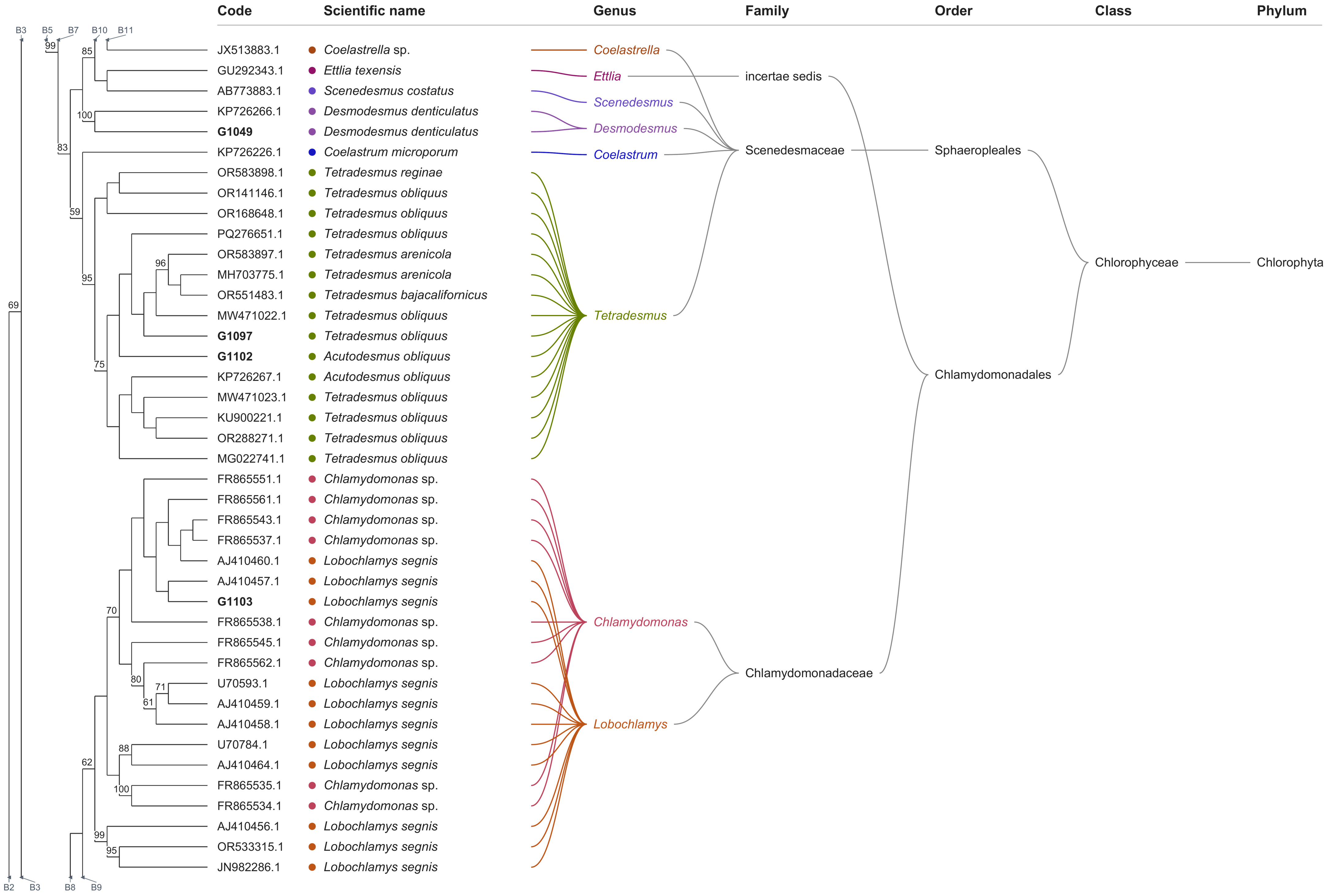


**Complete tree - part 2 of 4**

Tips 42-82 of 148

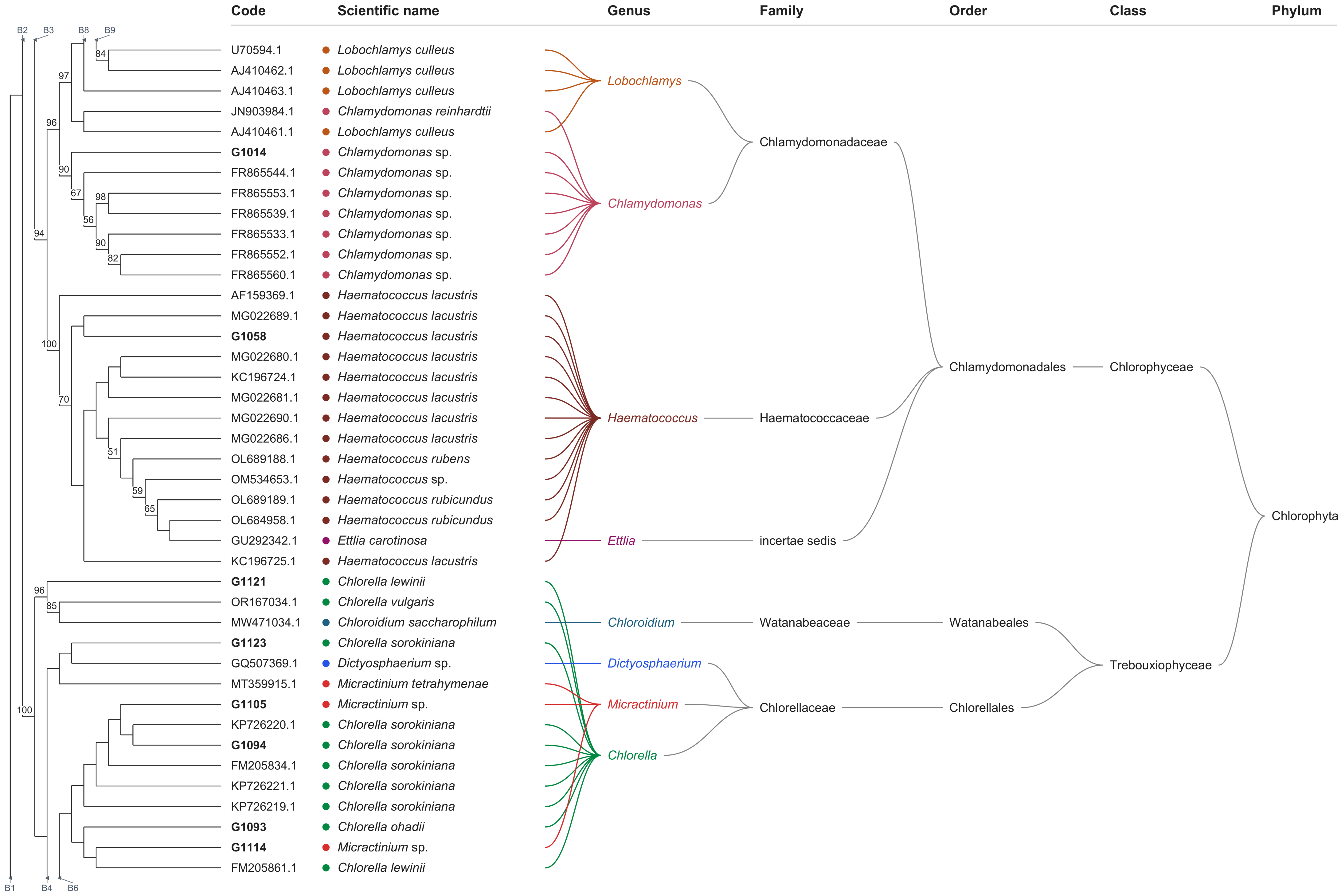


**Complete tree - part 3 of 4**

Tips 83-123 of 148

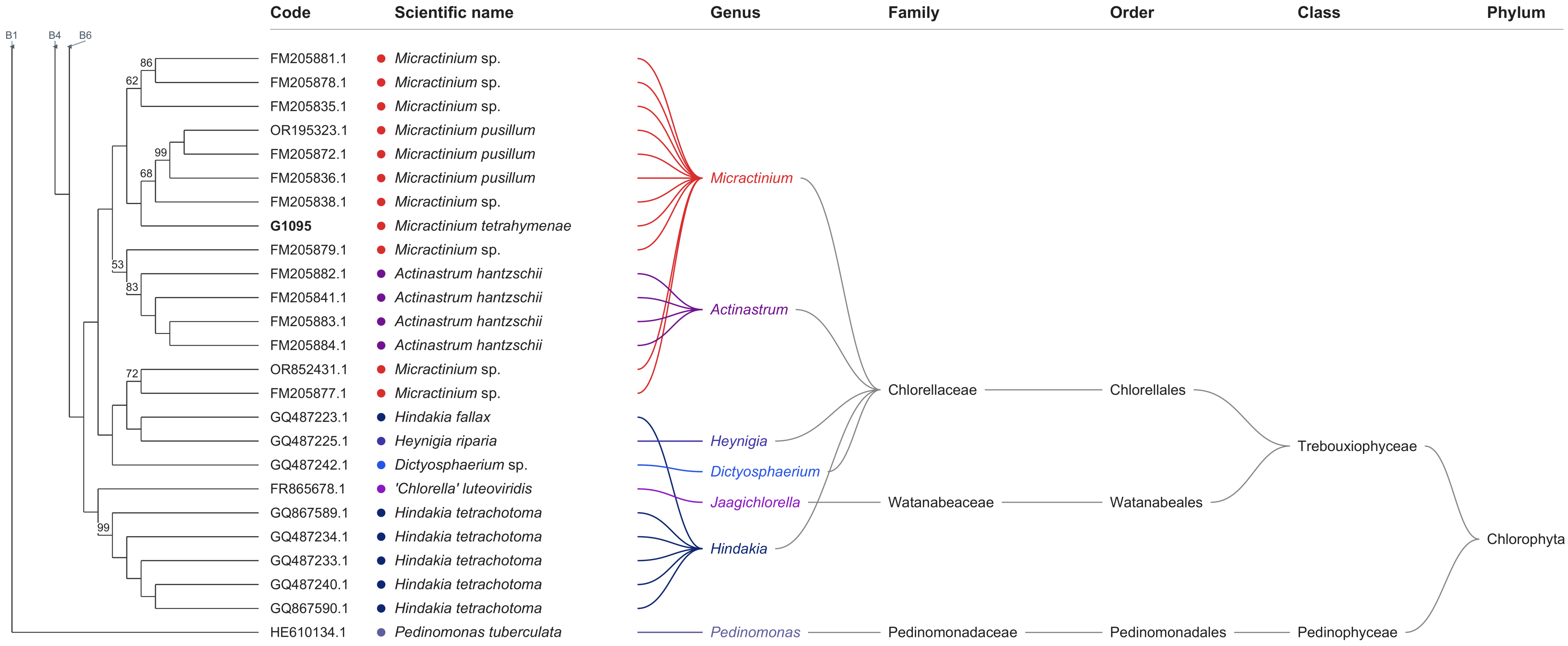


**Complete tree - part 4 of 4**

Tips 124-148 of 148

**Figure S7:** Fully expanded 18S phylogeny and reference-group key. The maximum-likelihood analysis in MEGA12 included 148 sequences comprising 20 study strains, 127 database references, and *Pedinomonas tuberculata* HE610134.1 as the outgroup. The analysis used the Tamura–Nei model with gamma-distributed rate variation (five categories) and invariant sites (TN93+G+I). Partial deletion at 95% site coverage retained 1,656 aligned 18S positions. Bootstrap support was estimated with 1,000 standard replicates. Branch spacing is schematic and does not represent evolutionary distance. All 148 tips are shown individually across four consecutive landscape pages. Table S4 lists all 127 reference accessions in the R01–R54 groups used in Figure 5. Read the tree pages from top to bottom. Branch coordinates are preserved across page boundaries, and matching B labels identify continuing branches; these are layout guides and not extra biological nodes. Taxonomic columns are repeated for the tips on each page. Bold codes identify the 20 study strains. Each genus has a distinct colour, shared by scientific-name circles and genus connectors. Bootstrap labels show percentages of at least 50, rounded to the nearest integer; zero-length-edge labels and colliding labels are omitted. Full numerical values are retained in the accompanying Newick tree. Scientific-name strings retain the study assignments and downloaded reference names; taxonomy notes document legacy reference names and species-specific classification. Identical or unresolved sequences should not be distinguished biologically from their schematic binary branching order. Taxonomic annotations do not independently validate the study identifications. *Acutodesmus*-labelled sequences are linked to *Tetradesmus* in the genus column. The retained reference label '*Chlorella*' *luteoviridis* (FR865678.1) is annotated as *Jaagichlorella* in Watanabeaceae/Watanabeales. *Ettlia* is shown as family *incertae sedis*; remaining legacy-name and species-specific taxonomy notes are supplied with the figure files.

**Table S4:** Reference key for Figure 5

127 reference sequences in 54 groups. n = references in each group. Read down the left block, then the right.

| Group | n | Accession | Scientific name |
|---|---|---|---|
| R01 | 2 | KF673369.1 | *Raphidocelis subcapitata* |
| | | KF144208.1 | *Monoraphidium* cf. *dybowskii* |
| R02 | 1 | KF673388.1 | *Ankistrodesmus fusiformis* |
| R03 | 1 | AJ300526.1 | *Monoraphidium neglectum* |
| R04 | 7 | MW471041.1 | *Ankistrodesmus* sp. |
| | | MW471043.1 | *Ankistrodesmus* sp. |
| | | MW471042.1 | *Ankistrodesmus* sp. |
| | | MW471046.1 | *Monoraphidium* sp. |
| | | MW471040.1 | *Ankistrodesmus* sp. |
| | | MW471047.1 | *Monoraphidium* sp. |
| | | MW471045.1 | *Ankistrodesmus* sp. |
| R05 | 2 | KP726271.1 | *Stauridium tetras* |
| | | AY780666.1 | *Stauridium tetras* |
| R06 | 1 | AY780665.1 | *Stauridium privum* |
| R07 | 2 | AY663030.1 | *Chlorotetraedron bitridens* |
| | | AY780669.1 | *Tetraedron caudatum* |
| R08 | 1 | AY780662.1 | *Pediastrum duplex* |
| R09 | 3 | OR791101.1 | *Hydrodictyon reticulatum* |
| | | OR791100.1 | *Hydrodictyon reticulatum* |
| | | AY780660.1 | *Hydrodictyon reticulatum* |
| R10 | 1 | OR178291.1 | *Pseudopediastrum boryanum* |
| R11 | 2 | MW471025.1 | *Desmodesmus abundans* |
| | | MK541740.1 | *Desmodesmus abundans* |
| R12 | 1 | MG022724.1 | *Desmodesmus abundans* |
| R13 | 1 | OR168646.1 | *Desmodesmus abundans* |
| R14 | 3 | MK541739.1 | *Desmodesmus abundans* |
| | | OR168645.1 | *Desmodesmus abundans* |
| | | KF673371.1 | *Desmodesmus abundans* |
| R15 | 1 | MW471024.1 | *Desmodesmus subspicatus* |
| R16 | 1 | FR865714.1 | *Desmodesmus perforatus* |
| R17 | 1 | MK541735.1 | *Desmodesmus armatus* |
| R18 | 2 | OR195326.1 | *Desmodesmus denticulatus* |
| | | OR195327.1 | *Desmodesmus denticulatus* |
| R19 | 1 | KM020087.1 | *Coelastrella* sp. |
| R20 | 1 | JX513883.1 | *Coelastrella* sp. |
| R21 | 2 | AB773883.1 | *Scenedesmus costatus* |
| | | GU292343.1 | *Ettlia texensis* |
| R22 | 1 | KP726266.1 | *Desmodesmus denticulatus* |
| R23 | 1 | KP726226.1 | *Coelastrum microporum* |
| R24 | 3 | OR168648.1 | *Tetradesmus obliquus* |
| | | OR583898.1 | *Tetradesmus reginae* |
| | | OR141146.1 | *Tetradesmus obliquus* |
| R25 | 1 | PQ276651.1 | *Tetradesmus obliquus* |
| R26 | 4 | OR551483.1 | *Tetradesmus bajacalifornicus* |
| | | MH703775.1 | *Tetradesmus arenicola* |
| | | OR583897.1 | *Tetradesmus arenicola* |
| | | MW471022.1 | *Tetradesmus obliquus* |
| R27 | 5 | MG022741.1 | *Tetradesmus obliquus* |
| | | KP726267.1 | *Acutodesmus obliquus* |
| | | MW471023.1 | *Tetradesmus obliquus* |
| | | KU900221.1 | *Tetradesmus obliquus* |
| | | OR288271.1 | *Tetradesmus obliquus* |
| R28 | 1 | FR865551.1 | *Chlamydomonas* sp. |
| R29 | 4 | FR865561.1 | *Chlamydomonas* sp. |
| | | AJ410460.1 | *Lobochlamys segnis* |
| | | FR865537.1 | *Chlamydomonas* sp. |
| | | FR865543.1 | *Chlamydomonas* sp. |
| R30 | 1 | AJ410457.1 | *Lobochlamys segnis* |
| R31 | 1 | FR865538.1 | *Chlamydomonas* sp. |
| R32 | 5 | AJ410459.1 | *Lobochlamys segnis* |
| | | U70593.1 | *Lobochlamys segnis* |
| | | AJ410458.1 | *Lobochlamys segnis* |
| | | FR865562.1 | *Chlamydomonas* sp. |
| | | FR865545.1 | *Chlamydomonas* sp. |
| R33 | 4 | AJ410464.1 | *Lobochlamys segnis* |
| | | U70784.1 | *Lobochlamys segnis* |
| | | FR865535.1 | *Chlamydomonas* sp. |
| | | FR865534.1 | *Chlamydomonas* sp. |
| R34 | 3 | JN982286.1 | *Lobochlamys segnis* |
| | | OR533315.1 | *Lobochlamys segnis* |
| | | AJ410456.1 | *Lobochlamys segnis* |
| R35 | 2 | AJ410462.1 | *Lobochlamys culleus* |
| | | U70594.1 | *Lobochlamys culleus* |
| R36 | 1 | AJ410463.1 | *Lobochlamys culleus* |
| R37 | 2 | JN903984.1 | *Chlamydomonas reinhardtii* |
| | | AJ410461.1 | *Lobochlamys culleus* |
| R38 | 6 | FR865560.1 | *Chlamydomonas* sp. |
| | | FR865552.1 | *Chlamydomonas* sp. |
| | | FR865533.1 | *Chlamydomonas* sp. |
| | | FR865539.1 | *Chlamydomonas* sp. |
| | | FR865553.1 | *Chlamydomonas* sp. |
| | | FR865544.1 | *Chlamydomonas* sp. |
| R39 | 1 | AF159369.1 | *Haematococcus lacustris* |
| R40 | 1 | MG022689.1 | *Haematococcus lacustris* |
| R41 | 11 | OL684958.1 | *Haematococcus rubicundus* |
| | | GU292342.1 | *Ettlia carotinosa* |
| | | OL689189.1 | *Haematococcus rubicundus* |
| | | OM534653.1 | *Haematococcus* sp. |
| | | OL689188.1 | *Haematococcus rubens* |
| | | MG022686.1 | *Haematococcus lacustris* |
| | | MG022690.1 | *Haematococcus lacustris* |
| | | MG022681.1 | *Haematococcus lacustris* |
| | | KC196724.1 | *Haematococcus lacustris* |
| | | MG022680.1 | *Haematococcus lacustris* |
| | | KC196725.1 | *Haematococcus lacustris* |
| R42 | 2 | MW471034.1 | *Chloroidium saccharophilum* |
| | | OR167034.1 | *Chlorella vulgaris* |
| R43 | 1 | GQ507369.1 | *Dictyosphaerium* sp. |
| R44 | 1 | MT359915.1 | *Micractinium tetrahymenae* |
| R45 | 1 | KP726220.1 | *Chlorella sorokiniana* |
| R46 | 1 | FM205834.1 | *Chlorella sorokiniana* |
| R47 | 1 | KP726221.1 | *Chlorella sorokiniana* |
| R48 | 1 | KP726219.1 | *Chlorella sorokiniana* |
| R49 | 1 | FM205861.1 | *Chlorella lewinii* |
| R50 | 3 | FM205881.1 | *Micractinium* sp. |
| | | FM205878.1 | *Micractinium* sp. |
| | | FM205835.1 | *Micractinium* sp. |
| R51 | 4 | FM205838.1 | *Micractinium* sp. |
| | | FM205836.1 | *Micractinium pusillum* |
| | | OR195323.1 | *Micractinium pusillum* |
| | | FM205872.1 | *Micractinium pusillum* |
| R52 | 5 | FM205884.1 | *Actinastrum hantzschii* |
| | | FM205883.1 | *Actinastrum hantzschii* |
| | | FM205841.1 | *Actinastrum hantzschii* |
| | | FM205882.1 | *Actinastrum hantzschii* |
| | | FM205879.1 | *Micractinium* sp. |
| R53 | 5 | FM205877.1 | *Micractinium* sp. |
| | | OR852431.1 | *Micractinium* sp. |
| | | GQ487225.1 | *Heynigia riparia* |
| | | GQ487223.1 | *Hindakia fallax* |
| | | GQ487242.1 | *Dictyosphaerium* sp. |
| R54 | 6 | GQ487240.1 | *Hindakia tetrachotoma* |
| | | GQ867590.1 | *Hindakia tetrachotoma* |
| | | GQ487233.1 | *Hindakia tetrachotoma* |
| | | GQ487234.1 | *Hindakia tetrachotoma* |
| | | GQ867589.1 | *Hindakia tetrachotoma* |
| | | FR865678.1 | *'Chlorella' luteoviridis* |

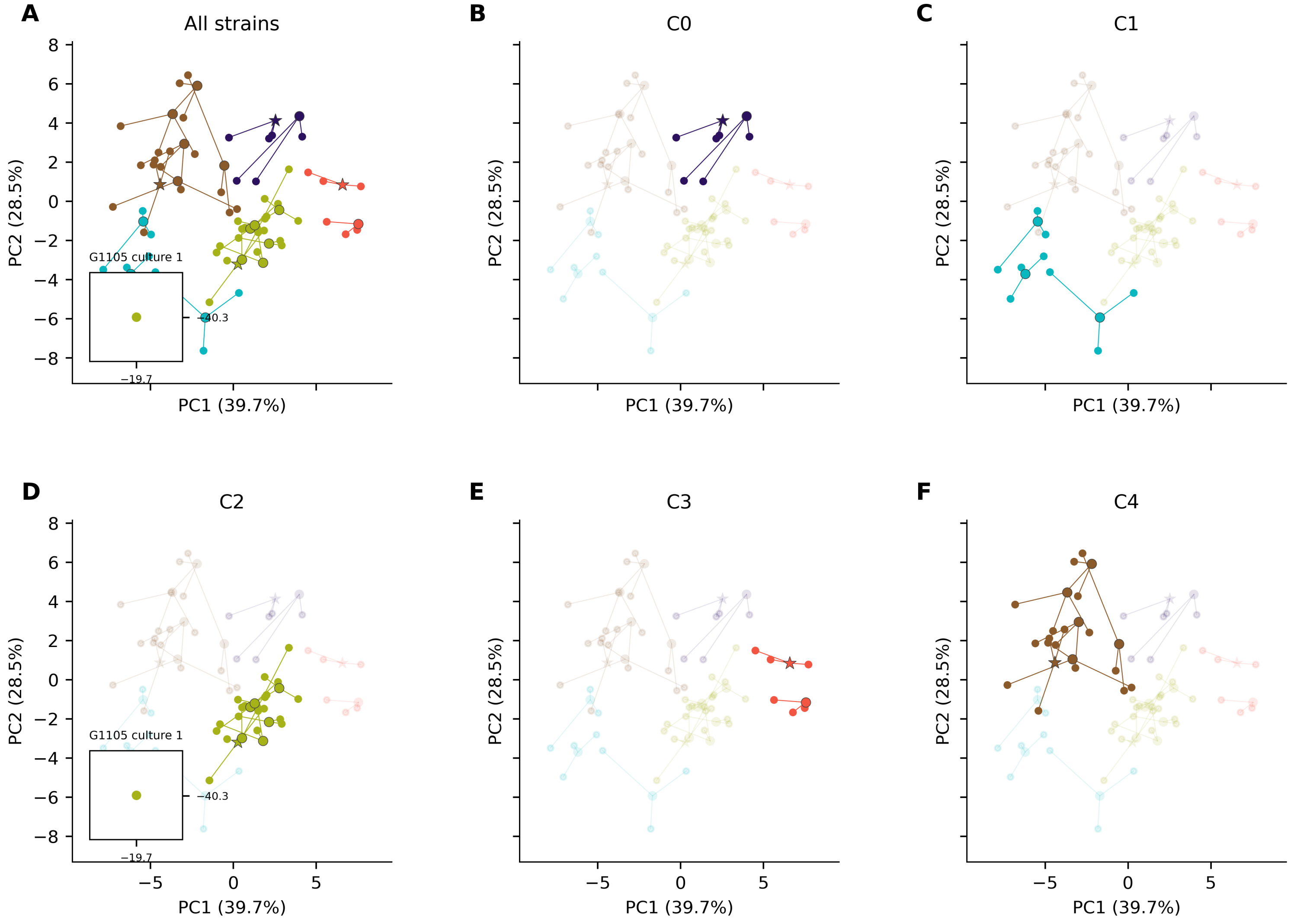


**Figure S8:** Biological replicate variation in $O_2$ exchange profiles. Large markers show strain means; small dots show 60 cultures (three per strain). Stars identify *Chlorella*; circles identify other strains. Lines connect cultures to their strain mean. Each culture's evolution and respiration profiles were separately peak-normalized and projected into the PCA fitted to strain means (Figure 7). Colors denote C0–C4 assignments (Figures 6 and 7). (A) All strains. (B–F) C0–C4 highlighted, respectively. PC1 and PC2 explain 39.7% and 28.5% of variation among means. Panels share axis limits. One G1105 culture appears in insets A and D (PC1 = −19.71; PC2 = −40.28). Its small respiration-normalization denominator (−2.68 μmol $O_2$ $L^{-1}$ $min^{-1}$) amplifies its atypical signed profile. It was retained; the cause remains unresolved. See Supplementary Methods.

**Table S5:** Agreement between physiological partitions and associations with phylogenetic distances and taxonomic assignments. Mantel results use the revised 18S tree in Figures 5 and S7.

| Analysis and distance | Strains / groups | Statistic | $R^2$ | P |
|---|---|---|---|---|
| **ARI, matched five-group partitions** | 20 / 5 | ARI = 0.726 | — | Not tested |
| **Mantel, all strains** | 20 / — | r = 0.103 | — | 0.24440 |
| **Genus PERMANOVA, 1 − r, all strains** | 20 / 12 | F[11,8] = 1.312 | 0.643 | 0.29233 |
| **Genus PERMANOVA, sqrt(1 − r), all strains** | 20 / 12 | F[11,8] = 1.056 | 0.592 | 0.41039 |
| **Genus PERMANOVA, 1 − r, replicated genera** | 12 / 4 | F[3,8] = 0.502 | 0.158 | 0.72755 |
| **Genus PERMANOVA, sqrt(1 − r), replicated genera** | 12 / 4 | F[3,8] = 0.530 | 0.166 | 0.89998 |
| **PERMDISP, sqrt(1 − r), replicated genera** | 12 / 4 | F[3,8] = 0.453 | — | 0.70831 |
| **Named-species PERMANOVA, 1 − r, exploratory** | 15 / 12 | F[11,3] = 1.302 | 0.827 | 0.40268 |
| **Named-species PERMANOVA, sqrt(1 − r), exploratory** | 15 / 12 | F[11,3] = 1.043 | 0.793 | 0.46696 |

Permutation tests used 99,999 randomizations and seed 0. Mantel tests are two-sided; PERMANOVA and PERMDISP use the upper tail. ARI is descriptive, without a P value. PERMANOVA reports pseudo-F; PERMDISP reports F. $R^2$ is the unadjusted fraction of distance-based sums of squares associated with the grouping, not prediction accuracy.

The all-strain genus model has 11 between-group and 8 residual degrees of freedom, including eight singleton genera. Replicated-genus models and PERMDISP have 3 and 8 degrees of freedom. Named-species models have 11 and 3, exclude sp. and cf. assignments, and are exploratory. Large $R^2$ values in these sparse models require caution.

PERMDISP uses bias-adjusted distances to spatial medians on the square-root-distance scale for the four replicated genera; it cannot assess singleton genera. P values are unadjusted, and sensitivity analyses are exploratory. Full procedures follow in the Supplementary Methods.

**Table S6:** Local comparison of the 20 study contigs against 127 retained public reference sequences. The strongest retained reference and the strongest named alternative were ranked by the bit score of their strongest high-scoring segment pair (HSP), then total query coverage, HSP identity and accession. Identity is for that HSP; query coverage is given as strongest-HSP/total coverage, where total coverage is the union of all HSP query intervals for that reference. Named alternatives exclude the assigned species and the *Tetradesmus obliquus*/*Acutodesmus obliquus* synonym pair and require a strongest HSP of at least 1,000 alignment columns. For genus-level or cf. assignments, a named reference is a comparison candidate. Reference names follow the saved records. The comparison is restricted to this finite reference pool and is not a new search of the complete nucleotide database. Full HSP lengths and scores are supplied with the data. Study accession entries are retained as recorded; G1058 is pending, and G1102 is a reference strain not included in the new GenBank submission (—).

| Strain INCCA code | Assigned name | Contig length (bp) | Strongest retained reference (accession; culture strain) | HSP identity (%) | HSP / total query coverage (%) | Named alternative (accession; identity %; HSP / total coverage %) | GenBank accession (this study) |
|---|---|---|---|---|---|---|---|
| **G1014** | *Chlamydomonas* sp. | 2166 | *Chlamydomonas* sp.; CCAP 11/138; FR865533.1 | 96.86 | 99.77/99.77 | *Lobochlamys culleus*; SAG 64.72; AJ410463.1; 98.16%; 77.79/77.79% | QB355095 |
| **G1044** | *Pediastrum duplex* | 2369 | *Hydrodictyon reticulatum*; CCAP 236-1A; OR791100.1 | 98.90 | 100.00/100.00 | *Hydrodictyon reticulatum*; CCAP 236-1A; OR791100.1; 98.90%; 100.00/100.00% | QB355096 |
| **G1049** | *Desmodesmus denticulatus* | 2353 | *Desmodesmus denticulatus*; KLL-G003; KP726266.1 | 99.07 | 100.00/100.00 | *Tetradesmus reginae*; CCAP:276/66; OR583898.1; 96.78%; 100.00/100.00% | QB355097 |
| **G1058** | *Haematococcus lacustris* | 2470 | *Haematococcus lacustris*; CCAP 34/13; MG022689.1 | 98.70 | 99.27/99.27 | *Ettlia carotinosa*; SAG 213-4; GU292342.1; 99.54%; 70.36/70.36% | QB360988 |
| **G1070** | *Stauridium tetras* | 2400 | *Stauridium tetras*; KLL-G013; KP726271.1 | 99.88 | 100.00/100.00 | *Hydrodictyon reticulatum*; CCAP 236-1A; OR791100.1; 94.25%; 100.00/100.00% | QB355098 |
| **G1074** | *Coelastrella* sp. | 2358 | *Coelastrella* sp.; SAG 2123; JX513883.1 | 98.90 | 99.96/99.96 | *Tetradesmus reginae*; CCAP:276/66; OR583898.1; 95.86%; 99.96/99.96% | QB355099 |
| **G1093** | *Chlorella ohadii* | 2413 | *Chlorella sorokiniana*; SAG 211-8k; FM205834.1 | 99.96 | 100.00/100.00 | *Chlorella sorokiniana*; SAG 211-8k; FM205834.1; 99.96%; 100.00/100.00% | QB355100 |
| **G1094** | *Chlorella sorokiniana* | 2410 | *Chlorella sorokiniana*; SAG 211-8k; FM205834.1 | 99.96 | 100.00/100.00 | *Micractinium tetrahymenae*; SAG 2587; MT359915.1; 94.53%; 98.63/98.63% | QB355101 |
| **G1095** | *Micractinium tetrahymenae* | 2450 | *Chlorella sorokiniana*; SAG 211-8k; FM205834.1 | 94.02 | 98.33/98.33 | *Chlorella sorokiniana*; SAG 211-8k; FM205834.1; 94.02%; 98.33/98.33% | QB355102 |

| Strain INCCA code | Assigned name | Contig length (bp) | Strongest retained reference (accession; culture strain) | HSP identity (%) | HSP / total query coverage (%) | Named alternative (accession; identity %; HSP / total coverage %) | GenBank accession (this study) |
|---|---|---|---|---|---|---|---|
| **G1096** | *Desmodesmus abundans* | 2305 | *Desmodesmus abundans*; CCAP 258/211; MW471025.1 | 99.96 | 100.00/100.00 | *Desmodesmus subspicatus*; CCAP 258/247; MW471024.1; 98.48%; 100.00/100.00% | QB355103 |
| **G1097** | *Tetradesmus obliquus* | 2358 | *Tetradesmus obliquus*; CCAP 276-6B; OR288271.1 | 100.00 | 100.00/100.00 | *Tetradesmus bajacalifornicus*; SAG:3.99; OR551483.1; 99.03%; 100.00/100.00% | QB355104 |
| **G1101** | *Chlorolobion braunii* | 2378 | *Ankistrodesmus* sp.; CCAP 202/8C; MW471041.1 | 99.75 | 100.00/100.00 | *Monoraphidium neglectum*; SAG 48.87; AJ300526.1; 99.54%; 73.00/73.00% | QB355105 |
| **G1102** | *Acutodesmus obliquus* | 2351 | *Tetradesmus obliquus*; CCAP 276/3C; MG022741.1 | 100.00 | 100.00/100.00 | *Tetradesmus bajacalifornicus*; SAG:3.99; OR551483.1; 99.07%; 100.00/100.00% | — |
| **G1103** | *Lobochlamys segnis* | 2347 | *Chlamydomonas* sp.; CCAP 11/167; FR865561.1 | 99.66 | 99.91/99.91 | *Lobochlamys culleus*; SAG 17.73; AJ410461.1; 99.20%; 74.22/74.22% | QB355106 |
| **G1104** | *Monoraphidium* cf. *dybowskii* | 2394 | *Ankistrodesmus* sp.; CCAP 202/9; MW471040.1 | 95.92 | 100.00/100.00 | *Monoraphidium neglectum*; SAG 48.87; AJ300526.1; 98.11%; 72.77/72.77% | QB355107 |
| **G1105** | *Micractinium* sp. | 2429 | *Chlorella sorokiniana*; SAG 211-8k; FM205834.1 | 96.96 | 100.00/100.00 | *Chlorella sorokiniana*; SAG 211-8k; FM205834.1; 96.96%; 100.00/100.00% | QB355108 |
| **G1107** | *Desmodesmus abundans* | 2305 | *Desmodesmus abundans*; CCAP 258/211; MW471025.1 | 100.00 | 100.00/100.00 | *Desmodesmus subspicatus*; CCAP 258/247; MW471024.1; 98.53%; 100.00/100.00% | QB355109 |
| **G1114** | *Micractinium* sp. | 2441 | *Chlorella sorokiniana*; SAG 211-8k; FM205834.1 | 97.02 | 100.00/100.00 | *Chlorella sorokiniana*; SAG 211-8k; FM205834.1; 97.02%; 100.00/100.00% | QB355110 |
| **G1121** | *Chlorella lewinii* | 2463 | *Chlorella vulgaris*; CCAP 211-51; OR167034.1 | 94.09 | 100.00/100.00 | *Chlorella vulgaris*; CCAP 211-51; OR167034.1; 94.09%; 100.00/100.00% | QB355111 |
| **G1123** | *Chlorella sorokiniana* | 1774 | *Chlorella sorokiniana*; KLL-G018; KP726220.1 | 100.00 | 100.00/100.00 | *Micractinium tetrahymenae*; SAG 2587; MT359915.1; 92.80%; 99.89/99.89% | QB355112 |

The working assignments are retained for traceability. G1044 has a stronger sequence match to *Hydrodictyon reticulatum* than to its morphology-based *Pediastrum duplex* assignment, which remains unresolved. G1095, G1101, G1104 and G1121 retain provisional species assignments, and the generic placement of G1105 and G1114 remains unresolved. The near-identity of the G1093 rDNA contig to *Chlorella sorokiniana* does not distinguish *C. ohadii* independently of culture provenance.

**Table S7:** ITS comparisons between strains sharing a nominal species assignment. Lengths follow the stated public reference annotations; they are reference-defined intervals rather than experimentally determined processing boundaries. Substitutions and indel bases are counted over the region available in both strains. G1123 has only 204 of the 229 reference-defined ITS2 bases represented in G1094, and G1102 has 245 of the 246 represented in G1097; missing sequence was excluded rather than counted as an indel. For *Desmodesmus*, ITS1 boundaries were transferred from *D. denticulatus* KP726266.1 using conserved flanking rRNA, whereas ITS2 boundaries came from the *D. abundans* record AJ400494.1. *Chlorella* boundaries follow FM205834.1 and *Tetradesmus* boundaries follow KP726267.1. Alternative *Chlorella* annotation conventions change the reported spacer lengths slightly but do not change the zero-difference result. CBC, compensatory base change: not assessed because no validated ITS2 secondary-structure comparison was performed. Full boundary coordinates, alignments and tied reference matches are included in the accompanying data. Values separated by a slash follow the strain order in column 2. Differences are substitutions/indel bases over the shared region; closest-reference entries are representative tied matches, each with 100% identity over its shared ITS region.

| Species assignment | Strains compared | ITS1 length (bp) | ITS2 length (bp) | ITS1 | ITS2 | ITS2 CBCs | Closest reference ITS (accession) |
|---|---|---|---|---|---|---|---|
| ***Chlorella sorokiniana*** | G1094 / G1123 | 272/272 | 229/204 (second partial) | 0/0 (272 bp) | 0/0 (204 bp) | Not assessed | ITS1: KP726220.1 (both); ITS2: FM205834.1 (both) |
| ***Desmodesmus abundans*** | G1096 / G1107 | 158/158 | 231/231 | 0/0 (158 bp) | 1/0 (231 bp) | Not assessed | ITS1: MW471025.1 (both); ITS2: OR168646.1 / MW471025.1 |
| ***Tetradesmus obliquus* (=*Acutodesmus obliquus*)** | G1097 / G1102 | 211/211 | 246/245 (second partial) | 0/0 (211 bp) | 1/0 (245 bp) | Not assessed | ITS1: KP726267.1 (both); ITS2: KP726267.1 / MG022741.1 |

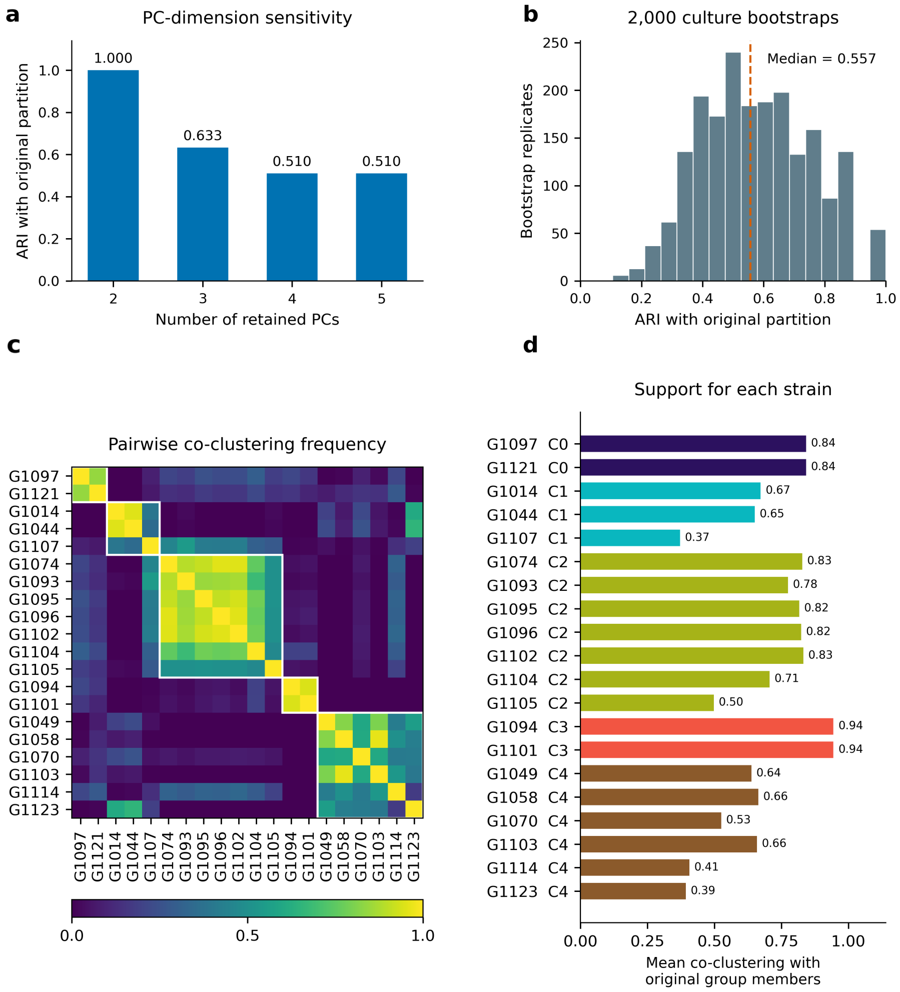


**Figure S9:** Sensitivity of the five-group physiological partition. (a) Adjusted Rand index (ARI) between the original PC1–PC2 partition and five-group k-means partitions calculated using the first two, three, four, or five PCs. (b) Distribution of ARI relative to the original partition across 2,000 within-strain culture bootstrap resamples; the dashed line marks the median. In each resample, three whole culture profiles were drawn with replacement from the three retained cultures of every strain, and the raw means, separate evolution and respiration normalization, variable standardization, PCA, and PC1–PC2 k-means clustering were recomputed. The group count remained fixed at five. All 2,000 resamples yielded valid five-group solutions and are included in every frequency denominator; none were discarded. (c) Frequency with which each pair of strains occurred in the same group. Strains are ordered by their original C0–C4 assignments; white boxes outline those groups, and diagonal values are one by definition. (d) For each strain, the mean pairwise co-clustering frequency with the other members of its original group, excluding itself. Bar colors denote the original groups. These frequencies are label-invariant and do not require matching numerical cluster labels. The bootstrap summarizes sensitivity to the three observed cultures per strain; it does not establish reproducibility in independently collected future cultures.

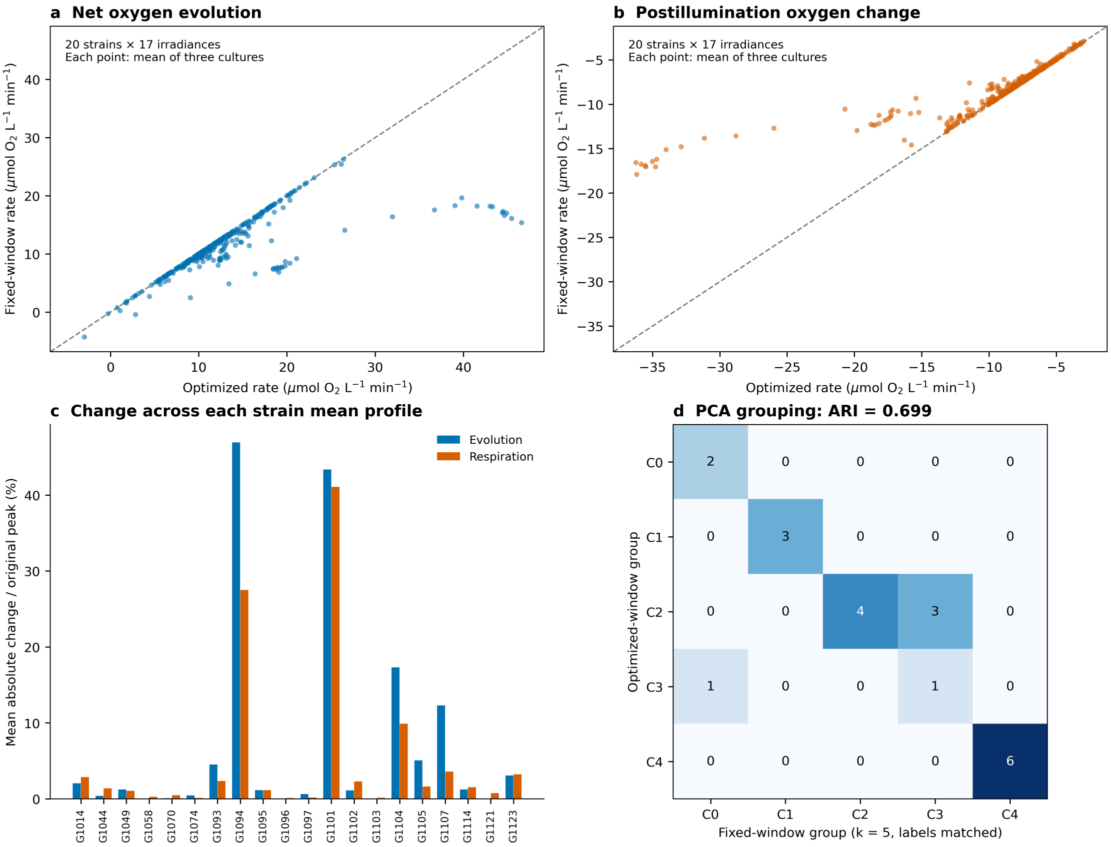


**Figure S10:** Sensitivity of $O_2$ exchange profiles to the rate-extraction window. The original maximum/minimum-slope procedure was compared with linear regression over the fixed central 15–45 s of each nominal 1-min light or postillumination dark interval in all 60 retained recordings. (a, b) Paired strain-mean rates for evolution and postillumination $O_2$ change, respectively; each panel contains 340 points (20 strains × 17 irradiances), each representing the mean of three cultures. Dashed lines indicate equality. Rates retain their signs, so negative values in b indicate net $O_2$ consumption. (c) Within each strain and process, the mean absolute difference between extraction methods across the 17 irradiances, expressed as a percentage of the absolute original optimized peak rate. (d) Cross-tabulation of the original and fixed-window PCA-based k-means partitions at k = 5. Cell values count strains; labels were matched by maximum membership overlap without changing assignments. Four strains changed group and ARI was 0.699. When k was selected independently, the fixed-window data favored four groups, with a mean silhouette coefficient of 0.452. No recordings, windows, or strains were omitted.

### Supplementary Methods

Vitamin Mix

A stock solution was prepared by dissolving thiamine hydrochloride (100 mg), biotin (1.5 mg), nicotinic acid (100 mg), vitamin $B_{12}$ (1 mg), and *p*-aminobenzoic acid (10 mg) in double-distilled water and adjusting the final volume to 100 mL. The solution was sterilized by filtration before addition to the culture medium. The stock was added to BG-11 at 1 mL per liter of medium.

Retained-reference sequence comparisons

The 20 retained Sanger consensus contigs were compared locally against the 127 public reference sequences retained for the 18S phylogeny using archived output from NCBI BLAST+ 2.17.0 (Camacho et al., 2009) (blastn, megablast task; E-value threshold 10; DUST and soft masking enabled). All retained query and reference sequences were verified as identical to the inputs of the archived comparison before the HSP results were filtered and reranked. The comparison was restricted to this finite reference pool and was not a new search against the complete nucleotide database. References were ranked by the bit score of their strongest high-scoring segment pair (HSP). For that HSP, we report identity, alignment length and query coverage; total query coverage was calculated separately as the union of query positions covered by all HSPs for the reference. Named alternatives excluded the assigned species and the *Acutodesmus obliquus*/*Tetradesmus obliquus* synonym pair and required a strongest HSP of at least 1,000 alignment columns. Ties were resolved by total query coverage, HSP identity and accession. Complete results within the retained pool are supplied in the accompanying data.

ITS boundary assignments and pairwise comparisons

ITS1 and ITS2 intervals were transferred from public INSDC feature annotations by local affine-gap alignment (Biopython 1.85; Cock et al., 2009; match +2, mismatch -3, gap opening -5, extension -2). Boundary references were FM205834.1 for the *Chlorella sorokiniana* pair, KP726266.1 for *Desmodesmus* ITS1, AJ400494.1 for *Desmodesmus* ITS2, and KP726267.1 for the *Tetradesmus obliquus* pair. The *Desmodesmus* ITS1 annotation was transferred from *D. denticulatus* using conserved flanking rRNA sequence; it is an operational homology-based boundary assignment. Reference-defined intervals were checked against their flanking rRNA sequence. Pairwise comparisons used global affine-gap alignments with the same scores and were restricted to the reference interval covered by both contigs. Substitutions and indel bases were counted separately. Missing terminal sequence was recorded as incomplete coverage, not a deletion. ITS2 secondary structures and compensatory base changes were not assessed.

Data processing and hierarchical clustering — detailed protocol (Figure 6)

**Step 1.** Use the final set of 60 independently grown cultures: three for each of the 20 strains. At each of the 17 irradiance levels, average the three $O_2$ evolution rates and, separately, the three respiration rates. This gives one mean value for each strain, irradiance level, and process. For G1103, the retained cultures were the three measured on 18 January 2026; the older recording from 24 December 2025 was not part of this final selection.

**Step 2.** Normalize each strain's profile separately for each process: divide every evolution value by that strain's own maximum mean evolution value and divide every respiration value by that strain's own most negative (largest-magnitude) mean respiration value. A value of 1 marks the strongest response for that process in that strain; values with the opposite sign to the normalization reference stay negative.

**Step 3.** Assemble these normalized mean values into a strains × irradiance-levels matrix for each process. This normalized matrix is the one displayed in the Figure 6 heatmap.

**Step 4.** For each strain, concatenate its 17 normalized evolution values and 17 normalized respiration values into one 34-variable vector.

**Step 5.** Check the 20 × 34 matrix for missing or non-numerical values. Every entry in the retained dataset was present and finite, so no imputation was performed. The same complete normalized matrix was used for the heatmap, its clustering, and the later distance-based comparisons.

**Step 6.** Run agglomerative hierarchical clustering on the 34-variable vectors in SciPy (scipy.cluster.hierarchy), using correlation distance (1 − Pearson's r) and average linkage.

**Step 7.** Compute the distance between two clusters as the mean of all pairwise distances between the strains they each contain. Apply optimal leaf ordering to the dendrogram for display — this reorders the leaves for readability and does not change the clustering hierarchy itself.

**Step 8.** Label every internal (merge) node with 1 minus its linkage distance — equivalently, the mean pairwise Pearson correlation between the two clusters being merged — and draw the dendrogram branches schematically (i.e., not scaled to clustering distance).

Principal component analysis and k-means clustering — detailed protocol (Figure 7)

Principal component analysis (PCA) and k-means clustering were used to compare the $O_2$ evolution and respiration profiles of the 20 green microalgal strains. The analysis was performed in Python using scikit-learn (Pedregosa et al., 2011), following the steps below:

**1. Retrieve the normalized mean profiles:** The normalized mean $O_2$ evolution and respiration values used in Figure 6 were retrieved. Each strain had been measured at 17 irradiance levels: 50, 100, 150, 200, 250, 300, 350, 400, 450, 500, 550, 600, 650, 700, 800, 1,000, and 2,000 µmol photons $m^{-2}$ $s^{-1}$. These normalized values described the relative response across the irradiance series. They were imported directly into the PCA workflow without repeating this normalization. For details how these normalized values were calculated see step 2 in "Data processing and hierarchical clustering — detailed protocol (Figure 6)".

**2.** Use the same retained cultures in every analysis: Assemble the strain means from the final 60-culture selection described above. Check that each strain has three retained cultures and that G-codes are used consistently across the input files. The strain-mean matrix is shared by Figures 6–8, the PCA reference space used for Figure S8, and the three analyses described below.

**3. Check measurement coverage and construct the data matrix:** Require 17 evolution values and 17 respiration values for every strain, in the same irradiance order. All 20 strains satisfied this requirement. Missing measurements should be resolved before analysis; they were not filled with replacement values in the present dataset.

The data retained was arranged into a matrix. Each row represented one strain. Each column represented one process at one measurement position, such as evolution at the first irradiance or respiration at the final irradiance.

The initial matrix therefore contained 34 variables per strain: 17 evolution values and 17 respiration values. Measurement positions were ordered consistently across all strains. For the 20 strains in this study, this corresponded to an initial matrix of 20 rows and 34 columns.

Each row represented a strain's mean physiological profile. Individual biological replicates were not entered as separate rows.

**4.** Check numerical completeness: Confirm that all $20 \times 34 = 680$ entries are finite **numerical values**. The retained matrix contained no missing entries and no wholly empty columns.

No mean imputation or removal of empty columns was therefore needed.

**5.** Check for **variables with negligible variation:** Calculate the variance of each column across the 20 strains and exclude a column if its variance is $\leq 10^{-12}$. Such a column would contain essentially no information for distinguishing strains. All 34 columns passed this check and were retained for PCA.

**6. Center and scale each retained variable:** Each remaining column was standardized across strains using scikit-learn's StandardScaler.

This standardization differed from the earlier normalization. Normalization expressed each strain's responses relative to its own strongest response. Standardization compared strains within each measurement column and placed the columns on a common scale. A negative standardized value indicated a response below the across-strain mean for that column. It did not, by itself, indicate negative $O_2$ evolution or respiration.

**7. Perform PCA and obtain the first two component scores:** PCA was applied to the standardized matrix using scikit-learn's PCA implementation. A strain's PC1 and PC2 scores provided its horizontal and vertical coordinates, respectively, on the two-dimensional PCA plot. The percentage of total variance explained by each component was reported with the corresponding axis.

**8. Use the unwhitened PC1 and PC2 scores for clustering:** K-means clustering used two values per strain: its PC1 score and its PC2 score. These scores were used without whitening, meaning that the two components were not rescaled to have equal variance. Consequently, clustering was performed on the strain positions in the PC1–PC2 plot, rather than directly on the original 34-variable profiles.

**9. Calculate candidate k-means solutions:** K-means clustering was evaluated for every integer value of k from 2 to 10, where k denotes the requested number of groups. Each candidate k was evaluated using 10 different initializations, with a fixed random seed of 0. For each k, scikit-learn retained the initialization with the lowest within-group sum of squared distances, also called inertia.

**10. Select the number of groups using the silhouette coefficient:** The retained solution for each candidate k was evaluated using the mean silhouette coefficient (Rousseeuw, 1987). Distances were calculated as ordinary straight-line, or Euclidean distances between strain positions in PC1–PC2 space. The silhouette values were averaged across strains to give one score for each candidate k. The solution with the highest mean score was selected.

**11. Display the selected groups and compare them with the heatmap:** Each strain was plotted at its PC1 and PC2 coordinates and labeled with its strain code. Colors indicated membership in the selected k-means groups. The same group assignments were transferred to the corresponding strains in the Figure 6 heatmap. Taxonomic identities were used to interpret the resulting patterns but were not included among the numerical variables used for PCA or k-means clustering. The selected solution had five groups and a mean silhouette coefficient of 0.508. PC1 and PC2 explained 39.7% and 28.5% of the variance, respectively. The final assignments place G1103 in C4; C0–C4 contain 2, 3, 7, 2, and 6 strains.

12. Project individual cultures without refitting the analysis: For each retained biological culture, divide its 17 evolution values by its own maximum evolution rate and its 17 respiration values by its own most negative respiration rate. Keep the same 34-column order as for the strain means. Apply the column means and standard deviations already fitted to the 20 strain means, then use that fitted PCA to obtain the culture's PC1 and PC2 coordinates. Do not fit a new scaler, PCA, or k-means model to the culture data. Color each culture according to the group assigned to its strain mean. Because strain means were calculated before peak normalization, a mean marker need not equal the average position of the three separately normalized culture markers. Figure S8 shows all 60 cultures. The extreme G1105 replicate is retained, with its coordinates shown in insets of panels A and D (PC1 = −19.71; PC2 = −40.28); only the main display limits omit that point.

**Shared preparation for ARI Mantel and PERMANOVA**

1. Define the unit being compared. Each of the 20 strains contributes one mean physiological profile. The three biological cultures for a strain contribute to that profile, but they are not counted as three separate taxonomic observations. The inputs therefore contain 20 observations, not 60 observations or 190 independent observations.

2. Use the current selected cultures. The dataset contains 60 cultures, with exactly three cultures per strain. For G1103, only the three cultures recorded on 18 January 2026 are retained. The earlier 24 December 2025 culture is excluded according to the selection used for Figures 6-8 and S8.

3. Assemble the complete physiological profile. Each row contains 34 numbers: 17 normalized $O_2$ evolution values followed by 17 normalized respiration values. Evolution and respiration were normalized separately within each strain as described above. All strains have all 34 values, so no missing-value replacement is needed. The order of the measurement positions is the same for every strain.

4. Keep the full physiological information for the distance analyses. Mantel and PERMANOVA use all 34 values. They do not use only the two axes drawn in the PCA plot, and they do not use the cluster labels. No additional feature-wise standardization across strains is applied before calculating these correlation distances.

5. Calculate one physiological distance for every pair of strains. First calculate Pearson's correlation between the two complete rows. Convert the correlation to a distance by subtracting it from 1. Arrange the distances in a 20 × 20 table, with the same strain names along the rows and columns.

**Adjusted Rand index Step by step**

Purpose. The adjusted Rand index, or ARI, measures agreement between two ways of grouping the same objects. Here the objects are the 20 strains, and the two groupings come from hierarchical clustering of the full physiological profiles and k-means clustering of the PCA scores. ARI was calculated following Hubert and Arabie (1985), using scikit-learn's adjusted_rand_score function.

1. Obtain the PCA-based groups. Start with the normalized 20 × 34 strain-mean matrix. Center and scale each column across strains, fit PCA, and run k-means on the unwhitened PC1 and PC2 scores as described above. Compare k = 2–10 using 10 initializations per k and random seed 0. The highest mean silhouette score selects five groups. Save each strain's identifier and its C0–C4 assignment.

2. Obtain the hierarchical groups. Calculate the 1 − Pearson correlation distances directly from the 34-value normalized profiles, without the extra column standardization used for PCA. Apply average-linkage hierarchical clustering. A dendrogram describes a sequence of merges; it does not automatically provide one fixed number of groups. Cut it at exactly five groups using SciPy's cut_tree function with n_clusters = 5. Five is chosen to match the PCA-based comparison, not because this step independently established five as the best hierarchical group count.

3. Calculate the index. Build a table whose rows are the k-means groups and whose columns are the hierarchical groups; each cell counts strains shared by that combination. Let A be the number of strain pairs together in k-means, B the number together in the hierarchy, and C the number together in both. A group of x strains contains $x(x - 1)/2$ pairs. Let T be the total number of pairs. Calculate expected shared pairs $E = A \times B / T$, then $ARI = (C - E) / [(A + B)/2 - E]$. Scikit-learn performs this calculation; the pair-count formula was also used as an independent check. No P value was calculated for the ARI.

4. Check the numerical result. In the present comparison, A = 41, B = 50, C = 36, and T = 190. These give ARI = 0.7263078, reported as 0.726. For comparison, 171 of the 190 pairs receive matching together-or-apart decisions before correcting for chance. The ARI value is the reported measure because it makes that chance adjustment.

5. Identify where the methods agree and differ. C0 and C1 are reproduced exactly by the hierarchical partition. The hierarchy combines all seven C2 strains and both C3 strains into one group. It divides the six C4 strains into G1070 alone and a group containing G1049, G1058, G1103, G1114, and G1123. Figure 6 displays H1–H5 alongside the colored PCA group

symbols so that these differences can be traced back to the heatmap. The complete strain assignments and contingency table are saved with the figure-generation program.

**Mantel test Step by step**

Purpose. The Mantel test (Mantel, 1967) asks whether pairs of strains that are more distant in the 18S tree also tend to be more different in their normalized $O_2$ exchange profiles.

1. Obtain the phylogenetic distances. Read the branch-length tree used for the updated Figure 5. For each pair of study strains, trace the path connecting their tips and add the branch lengths along that path. This is the patristic distance, expressed in the branch-length units of the tree. Bootstrap percentages and distances measured from the printed drawing are not used.

2. Select matching strains. Locate each of the 20 study strains exactly once in the tree. Reference sequences and the outgroup remain part of the estimated tree, but only the study strains have matching physiological observations and therefore enter this comparison. No additional observation is created from a reference sequence.

3. Calculate the observed association. Calculate Pearson's correlation between the 190 phylogenetic distances and the corresponding 190 physiological distances. This number is the Mantel r.

4. Create a randomized comparison. Randomly rearrange the strain labels in the physiological matrix. Move its rows and matching columns together. This breaks the match between physiological and phylogenetic identities while retaining the original physiological distance relationships. Individual pairwise entries are never shuffled independently.

5. Repeat the randomization. Calculate a new matrix correlation after each of 99,999 random label permutations. Use NumPy's default random-number generator with seed 0 so that running the script again reproduces the same randomization sequence.

6. Calculate a two-sided P value. Count the randomized correlations whose absolute value is at least as large as the absolute observed correlation. Add one to this count and divide by 100,000, which is the number of randomizations plus the original observation. The ordinary Pearson correlation P value is not used, because it would incorrectly treat the pairwise entries as independent observations.

**PERMANOVA Step by step**

Purpose. PERMANOVA (Anderson, 2001) asks whether strains assigned to the same genus tend to have more similar physiological profiles than would be expected if genus labels were randomly reassigned among the sampled strains.

1. Start with the physiological matrix above. Use the same 20 × 20 table of 1 − Pearson correlation distances. The phylogenetic distance matrix is not required for PERMANOVA. The grouping variable comes from the taxonomic assignments in Table 1.

Step 2. Prepare the genus labels. Give each strain one genus assignment. Harmonize the supplied name *Acutodesmus obliquus* for G1102 to *Tetradesmus obliquus* so that this taxonomic synonym does not create an artificial extra genus. This grouping step does not change the supplied culture name reported in Table 1. The resulting dataset contains 12 genera: *Chlorella* has four strains; *Desmodesmus* and *Micractinium* have three each; *Tetradesmus* has two; the remaining eight genera have one strain each.

3. Calculate the sums of squares. Following Anderson (2001), the total sum of squares is the sum of squared pairwise distances divided by the number of strains; the within-genus sum of squares is the corresponding quantity calculated within each genus and summed across genera. Singleton genera contribute zero to the within-genus sum of squares.

4. Calculate the part associated with genus. Subtract the within-genus sum of squares from the total sum of squares. Divide this between-genus quantity by the total to obtain the unadjusted $R^2$. $R^2$ is the fraction of the fitted distance-based sums of squares associated with the supplied grouping.

5. Calculate the pseudo-F statistic. Divide the between-genus sum of squares by the number of genera minus one. Divide the within-genus sum of squares by the number of strains minus the number of genera. The ratio of these two quantities is pseudo-F. The full dataset therefore has 11 between-genus and 8 within-genus degrees of freedom.

6. Assess the observed grouping by randomization. Randomly reassign the existing genus labels among the 20 strains while keeping the number of strains assigned to each genus unchanged. Recalculate pseudo-F for each randomized assignment. Repeat 99,999 times using NumPy's default random-number generator with seed 0. The P value is the number of randomized pseudo-F values at least as large as the observed value, plus one, divided by 100,000. The statistical unit being permuted is the complete strain.

7. Check sensitivity to singleton genera. Repeat the test using only the four genera with at least two strains. This retains 12 strains: four *Chlorella*, three *Desmodesmus*, three *Micractinium*, and two *Tetradesmus*. The restricted model has 3 between-genus and 8 within-genus degrees

of freedom. It addresses the subset of replicated genera and is not a replacement for the all-strain result.

8. Check the distance geometry. Principal-coordinate analysis of the original 1 − correlation matrix produced 11 negative eigenvalues below $-10^{-10}$, with a minimum eigenvalue of −0.50196. This means that those distances cannot be represented exactly as ordinary Euclidean distances. The primary PERMANOVA retains the original Figure 6 distance. As an explicit sensitivity check, repeat both the full and replicated-genus tests after taking the square root of every distance. Square-root correlation distances are Euclidean for these complete profiles, and no negative eigenvalue larger than numerical rounding remained. This transformation changes the sums of squares and is therefore reported as a separate analysis.

9. Examine differences in within-genus spread. On the square-root-distance scale, calculate the spatial median of each of the four replicated genera. Calculate the Euclidean distance of each strain to its own genus median, retaining every dimension. The implementation uses an exact Euclidean representation obtained by centering each 34-value profile on its own mean, scaling it to unit length, and dividing by sqrt(2). This representation reproduces the square-root correlation distances without reducing them to two PCA axes.

10. Account for the small, unequal genus sample sizes. Multiply each distance to its genus median by sqrt[n_g/(n_g-1)], where n_g is the number of strains in that genus. Use a one-factor ANOVA on these adjusted distances to obtain the dispersion F statistic. Subtract each genus's mean adjusted distance to obtain model residuals. Following the residual-permutation procedure in vegan's permutest.betadisper, permute these residuals among strains, refit the group model, and calculate the randomized F statistic. Use 99,999 permutations and seed 0. The medians are not recalculated during this residual randomization. This supporting test is called PERMDISP (Anderson, 2006). It concerns the square-root-distance analysis and the four replicated genera only. It cannot assess dispersion in the eight singleton genera, and a nonsignificant result does not establish equal dispersion.

11. Keep the species-level comparison exploratory. Exclude *Chlamydomonas* sp. G1014, *Coelastrella* sp. G1074, *Monoraphidium* cf. *dybowskii* G1104, and *Micractinium* sp. G1105 and G1114 from the strict named-species comparison. After synonym harmonization, the remaining 15 strains represent 12 named species. Only *Chlorella sorokiniana*, *Desmodesmus abundans* and *Tetradesmus obliquus* have two strains each. The other nine species have one strain each. Both distance versions are reported as exploratory sensitivity analyses, with only three residual degrees of freedom.

**Reproducing the calculations**

The supplied rebuild_multivariate.py program regenerates the normalized strain means, PCA and k-means results, hierarchical comparison, Mantel test, PERMANOVA and PERMDISP tests, PC-dimension sensitivity, culture bootstrap, and fixed-window sensitivity. The accompanying inputs folder contains the complete 60-culture rate matrix, current 18S branch-length tree, taxonomic assignments, and retained-culture manifest. The defaults for permutation tests are 99,999 permutations and seed 0; the culture bootstrap uses 2,000 resamples and seed 20261017. The render_figures.py program regenerates Figures 6–8, S8, and S10, while the bootstrap analysis writes Figure S9. The numeric CSV and JSON outputs record all assignments and settings.

The recomputed analyses used Python 3.12.14, NumPy 2.3.5, pandas 3.0.1, SciPy 1.15.3, scikit-learn 1.6.1, and Matplotlib 3.10.1. The supplied calculations implement the stated formulas directly; neither vegan nor scikit-bio was executed for these results. Independent calculations verified the patristic distances, the permutation results, the PERMANOVA sums of squares, and the dispersion diagnostic. All reported P values are unadjusted; the additional subsets, distance transformations, and dispersion checks are exploratory sensitivity analyses. A nonsignificant result is not proof that the underlying biological association or dispersion difference is absent.

Original-unit comparisons and culture-bootstrap sensitivity

The retained input consisted of 60 complete culture profiles (20 strains × 3 cultures × 34 rates), including the three 18 January 2026 G1103 cultures and excluding the previously omitted 24 December 2025 G1103 culture. Each strain had exactly three retained cultures with finite values for all 34 rates. Strain means, separate process normalization, variable standardization, PCA, and k-means were recomputed from this retained dataset. All 34 variables passed the variance check. Selection across k = 2–10 gave a five-group solution, with group sizes 2, 3, 7, 2, and 6 and silhouette coefficient 0.508467190. Numerical C0–C4 labels were matched by maximum membership overlap solely for visual continuity; the fitted memberships and silhouette were unchanged by this relabeling.

For PC-dimension sensitivity, a single PCA was fitted to the 34 standardized normalized strain-mean variables and k-means with five groups was fitted separately to the first two, three, four, or five score columns. For every fit, k-means used k-means++ initialization, 10 initializations, and random_state = 0 (scikit-learn 1.6.1). The group count was not reselected in these analyses. The ARI is invariant to arbitrary cluster labels. Numerical labels in the supplementary membership CSV were aligned to the original labels by maximum membership overlap using the Hungarian assignment algorithm solely for readability; this did not change any membership or ARI.

For the bootstrap, a NumPy default_rng generator with seed 20261017 generated 2,000 resamples. Within each resample and strain, three culture indices were sampled independently with replacement from the three retained cultures, preserving the entire 34-variable profile of each sampled culture. The three sampled profiles were averaged on the original signed-rate scale. The 17 mean evolution rates were divided by their maximum, and the 17 mean respiration slopes by their minimum. Variables with across-strain variance $\leq 10^{-12}$ would have been omitted, following the original pipeline; all 34 variables were retained in every resample. StandardScaler then centered each variable and scaled it to unit population standard deviation across the 20 strain means. PCA was refitted, followed by five-group k-means on the first two PC scores with the same initialization settings as above. All 2,000 resamples had finite normalization denominators and five nonempty clusters; none failed, were replaced, or were omitted. The normalization, standardization, and PCA were recomputed in each resample rather than projecting bootstrap profiles into fixed original axes.

Pairwise co-clustering frequency for strains i and j was the number of resamples assigning them to the same group divided by 2,000. For strain i, original-group support was the arithmetic mean of those frequencies over the other strains sharing i's original group, excluding i itself. This statistic can remain high when a group merges with additional strains and must not be interpreted as exact recovery of the original group. Separate CSV columns report the proportion of resamples retaining all original group members together and the stricter proportion recovering exactly the original membership without additional strains. Bootstrap ARIs were calculated against the complete original 20-strain partition. The reported 2.5th and 97.5th percentiles are empirical quantiles of the bootstrap ARI distribution, not confidence limits on a population parameter. All probabilities use all 2,000 resamples as their denominator. The analysis is conditional on the selected five-group model and the three retained cultures per strain.

The Figure 9 comparison retained all 17 irradiances and all three cultures for each of the six strains. For each pair/process/irradiance, the released table reports the two strain means, observed ranges, their signed mean difference, and whether the ranges were completely separated. Complete separation was evaluated by comparing all three observations from one strain with all three from the other; the nine cross-comparisons are dependent descriptions, not nine independent replicates. No significance test was applied to these selected peak differences, and the 17 irradiance steps were not treated as independent observations.

Fixed-window rate-extraction sensitivity

Fixed-window sensitivity analysis. All 60 recordings retained for the primary analysis were identified by an explicit filename manifest and verified against their recorded SHA-256 hashes. No recordings were excluded or substituted. The original code assigned light intervals to 1–2, 3–4, …, 33–34 min and the corresponding postillumination dark intervals to 2–3, 4–5, …, 34–35 min. The initial 0–1-min dark interval was not included in the irradiance response profiles. These are nominal intervals relative to recording time; no new estimate of the actual switching times or temporal realignment was introduced.

The original optimized rates were reproduced from channel 1 $O_2$ concentrations in µmol $L^{-1}$. Duplicate time points were handled as in the original pipeline, and concentration at 1 min was subtracted as a vertical baseline adjustment. Linear interpolation onto a 1.2-s grid provided 26 samples spanning each 30-s window. Original candidate starts ranged from −15 to +15 s relative to each nominal transition; the selected window maximized the light-associated slope or minimized the dark-associated slope. In the sensitivity analysis, every window instead began 15 s after the nominal transition and ended 45 s after it. This fixed window was the final candidate in the original search range and lay entirely within the nominal interval. All 2,040 fixed windows were covered by recorded observations; the largest gap between observations bracketing a fixed window was 3.05 s. No extrapolation or missing-profile imputation was used.

The reproduced optimized slopes agreed with the archived individual rates to within $1.5 \times 10^{-12}$ µmol $O_2$ $L^{-1}$ $min^{-1}$. The fixed-window regression slopes were independently checked with the centered-sum formula for ordinary least squares, with a maximum discrepancy below $3.4 \times 10^{-13}$ in the same units. The original selected windows crossed a nominal transition in 39 of 2,040 cases. Comparisons between extraction methods were paired by recording, process, and irradiance. Rates retained their signs; negative dark-phase slopes represented net $O_2$ consumption.

For each strain and process, rates were averaged across the same three cultures at each irradiance. Mean absolute change was calculated across the 17 irradiances and divided by the absolute optimized peak rate of that strain and process to express it as a percentage. Across-strain summaries therefore comprised 20 values per process. Mean profiles were then normalized separately to their own evolution maxima and respiration minima for each extraction method. PCA, variable standardization, and k-means clustering were refitted using the original settings. Cluster agreement was evaluated both at a fixed k = 5 and after independently selecting k from 2–10 by the mean silhouette coefficient. Numeric labels in the five-group comparison were matched to the primary groups by maximum membership overlap; this relabeling did not change assignments.

For each pair sharing a species assignment, the direction of the difference between strain means was compared between extraction methods at all 17 irradiances for both processes. Culture-range separation was also recorded when every observed rate for one strain exceeded every rate for the other; overlapping ranges were not described as consistently ordered. These are descriptive sensitivity comparisons, not tests treating irradiances or interpolated points as independent replicates.
